# Epigenomic regulatory programs reveal the latent heterogeneity of complex disease

**DOI:** 10.64898/2026.05.01.722123

**Authors:** Woo Jun Shim, Shaine Chenxin Bao, Chris Siu Yeung Chow, Sophie Shen, Zachary Riedlshah, Qiongyi Zhao, Jian Zeng, Sonia Shah, Dalia Mizikovsky, Mikael Boden, Nathan J. Palpant

## Abstract

Complex diseases exhibit substantial biological and clinical heterogeneity despite shared diagnoses. Here, we show that disease-associated variants are organised through recurrent epigenomic programs that capture coordinated regulatory activity across human tissues and cell states. These programs, termed Epigenomically Co-modulated programs (EpiCops), provide a disease-agnostic representation to partition disease-associated variants without phenotypic or disease-specific priors, revealing distinct tissue, cellular and chromatin contributions to disease risk. Applied to type 2 diabetes, EpiCops resolve risk variants into distinct regulatory profiles characterised by contrasting tissue contributions and epigenomic features with divergent cardiometabolic risks of myocardial infarction and non-alcoholic fatty liver disease. Together, EpiCops provide a generalisable epigenomic framework for linking inherited disease risk to cellular regulatory context and reveal how distinct regulatory architectures underlie biological and clinical heterogeneity of complex disease.

## Introduction

Complex diseases are commonly defined by shared clinical features, yet individuals with the same diagnosis can differ substantially in pathophysiology, disease progression and therapeutic response, highlighting biological heterogeneity that is not captured by diagnostic labels alone^1, 2^. Most genetic approaches to study complex disease nevertheless assume that disease cases represent a relatively homogeneous group in which genetic liability is additive across variants^3, 4^. Genome-wide association studies (GWAS) identify disease-associated variants across the case cohort and estimate their average effects while polygenic scores (PGS) sum these effects into a single measure of inherited susceptibility^5–7^. Although these approaches have been highly effective for total risk prediction, aggregate models provide limited information on biological subtypes driven by distinct biological mechanisms. This limitation is particularly important when variants associated with the same disease act through different pathways or cellular contexts, as divergent effects across subtypes may be diluted^5^ or cancelled^7^ when aggregated across variants.

Resolving this heterogeneity requires understanding the biological contexts through which disease-associated variants act. Most variants associated with complex disease lie within non-coding regulatory regions, with genetic risk enriched in regulatory elements that are active in specific tissues and cell types^8^. These elements do not act independently, but contribute to coordinated regulatory states through patterns of chromatin accessibility and histone modification that define cellular identity and function^9–11^. Genetic variants contributing to the same disease may therefore act through different combinations of regulatory programs across tissues and cell states, providing a potential basis for resolving different biological pathways that converge on a common clinical diagnosis.

Existing approaches to resolve disease heterogeneity commonly group individuals or disease-associated variants according to clinical variables^12^, molecular features^13^, pathways or auxiliary trait associations^14–19^, and have revealed meaningful disease subtypes. However, approaches that assign genetic effects to known pathways and cellular processes often depend on existing biological knowledge, including curated gene sets, functional annotations, predefined tissues or cell types, and selected disease-relevant phenotypes. Such dependencies may constrain mechanistic insights to those defined by the input framework. Similarly, subtyping based on downstream molecular or clinical features can capture disease progression and treatment effects as well as inherited biology, making it difficult to disentangle the upstream genetic mechanisms from downstream disease manifestations.

Epigenomic information provides a complementary layer for interpreting disease-associated variation by linking genetic loci to regulatory elements, chromatin states and disease-relevant cell types^9–11^. Consortium-scale epigenomic atlases have now mapped these features across diverse human tissues and cell states^20–23^, providing an extensive resource for characterising the regulatory contexts in which genetic variants may act. However, these data have been largely used to annotate disease variants retrospectively or test enrichment within predefined regulatory features, tissues or cell types^24, 25^. Such analyses do not fully capture how regulatory activity is organised into shared programs across the genome or whether disease-associated variants participate in these programs. This distinction is important for complex diseases, where variants contributing to the same diagnosis may act through different combinations of regulatory contexts across diverse cell types to govern distinct underlying disease subtypes and mechanisms.

We hypothesised that genome-wide epigenomic activity is organised into recurrent regulatory programs that capture coordinated patterns of chromatin activity across tissues and cell states. Under this model, variants contributing to the same disease can be partitioned according to the regulatory programs through which they act, rather than their association with pre-defined clinical phenotype. Distinct subsets of variants contributing to the same disease may therefore map to different combinations of tissues, cell states and chromatin features, revealing biological heterogeneity obscured when genetic effects are aggregated. Importantly, because these programs are learned independently of disease, they provide a common framework for resolving regulatory heterogeneity across diseases without requiring phenotypic or disease-specific priors for variant partitioning.

Here, we develop Epigenomically Co-modulated programs (EpiCops), a disease-agnostic epigenomic framework that learns recurrent regulatory programs from consortium-scale maps of chromatin accessibility and histone modifications across human tissues and cell states^20, 21^. These regulatory programs capture coordinated patterns of epigenomic activity across the genome and provide a disease-independent representation for organising genetic variation by regulatory contexts. We show that EpiCops recover biological structure across genomic loci from diverse contexts and partition disease-associated variants into regulatory profiles defined by different tissue, cellular and chromatin activities. In type 2 diabetes, these programs resolve genetic variants into biologically distinct subgroups with contrasting tissue distributions and epigenomic features and are associated with divergent cardiometabolic risks and disease outcomes. Together, these findings establish EpiCops as a generalisable epigenomic framework for resolving regulatory heterogeneity in complex disease by partitioning genetic risk into regulatory programs with distinct tissue and cellular mechanisms.

## Results

### EpiCops are recurrent epigenomic programs extracted from a consortium-scale epigenomic atlas

To construct a disease-agnostic representation of genomic regulatory activity, we integrated consortium-scale chromatin accessibility and histone modification profiles from EpiMap, spanning 6,181 epigenomic samples across 833 tissues and cell states^20^. Collectively, these datasets contained 4.61 billion chromatin peaks covering approximately 85% of the human genome (**Supplementary Fig. 1a-c**). The peaks were enriched at regulatory elements including transcription start sites^26^, FANTOM5 enhancers^27^, and long non-coding RNA loci^28^ (**Supplementary Fig. 1d**), suggesting their preferential localisation at regulatory elements in the genome. Using peak representations across non-overlapping 1-kb genomic windows, restricted Boltzmann machines (RBMs)^29^ were trained independently for each chromatin mark to learn recurrent patterns of their activity across tissues and cell states as latent representations (**Fig. 1a**). Following the model optimisation (**Supplementary Fig. 2a**), we selected the number of learned features that minimised the reconstruction error in held-out data (**Supplementary Fig. 2b**).

**Figure 1.**
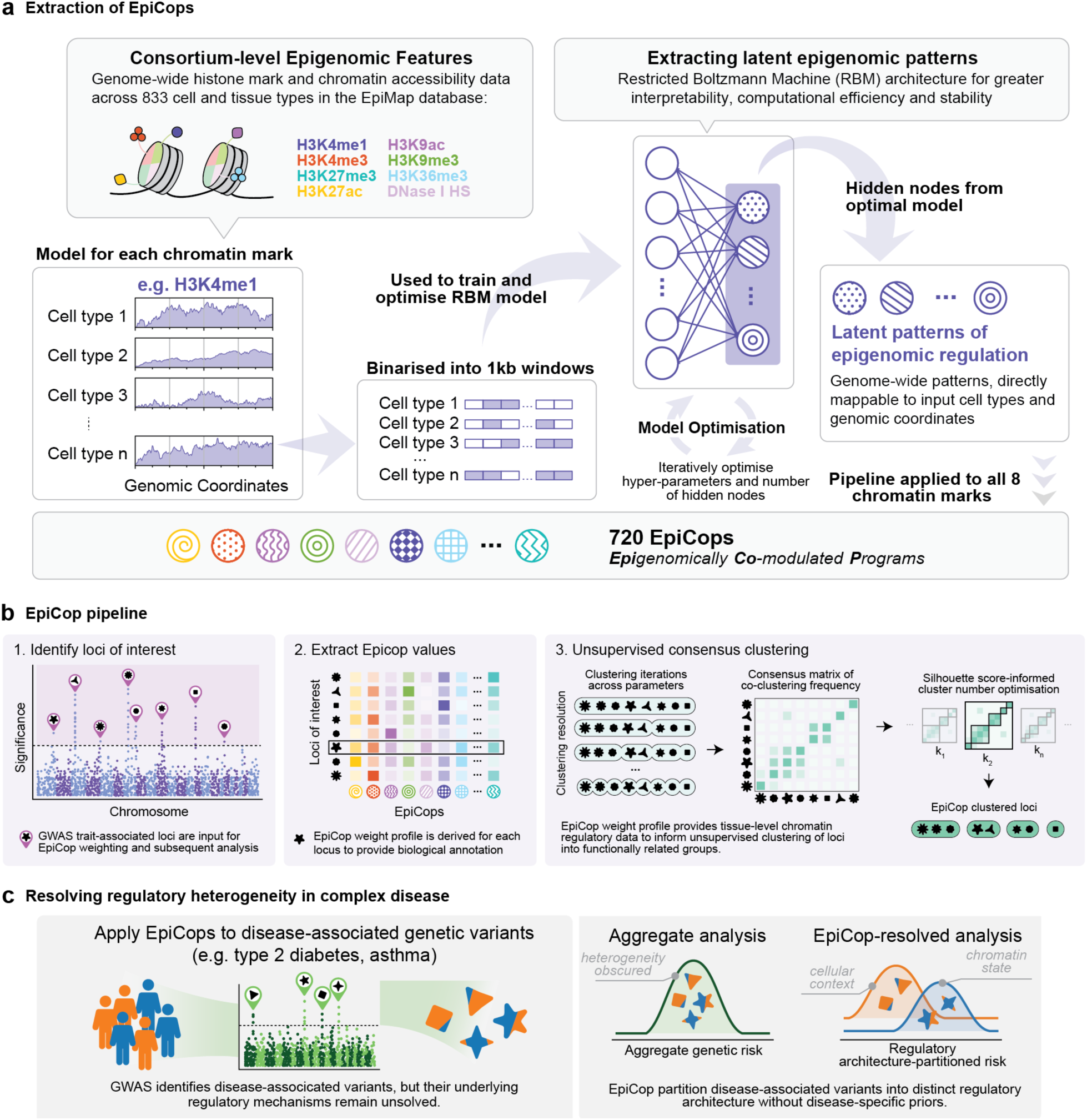
Overview of the EpiCop framework. **a.** Extraction of EpiCops. Consortium-scale epigenomic profiles across 833 EpiMap cell and tissue samples were analysed using chromatin mark-specific restricted Boltzmann machine models to identify latent regulatory patterns of coordinated epigenomic activity. Across eight chromatin marks, 720 EpiCops were identified. **b.** EpiCop pipeline. Genomic loci of interest are mapped to EpiCop profiles and grouped by a unsupervised consensus clustering approach to identify loci with shared tissue, cellular and chromatin regulatory patterns. **c.** Application of EpiCops to dissect heterogeneity of complex disease. Disease-associated variants are partitioned into subgroups with distinct cell-type specific epigenomic structures, enabling functional characterisation of disease subtypes and construction of subtype-specific polygenic risk scores.

Across the eight chromatin marks, the optimised models yielded 720 latent regulatory patterns, which we term Epigenomically Co-modulated Programs (EpiCops). Defined by the vector of weights across EpiMap samples, each EpiCop captures a coordinated pattern of chromatin activity, enabling direct interpretation of its tissue and cellular context. EpiCops provide a common set of regulatory dimensions onto which genomic loci of interest can be mapped. The analytical pipeline begins with an identification of loci of interest, such as disease-associated variants, from which EpiCop values are extracted. These EpiCop profiles are then used in an unsupervised consensus clustering framework^30^ to group loci according to shared patterns of epigenomic activity across tissues and cell states (**Fig. 1b**). Applied to complex disease GWAS data, this approach provides a disease-agnostic means to group genetic variants by distinct regulatory programs to subsequently examine their tissue, cellular and phenotypic associations (**Fig. 1c**).

### EpiCops capture distinct and reproducible regulatory programs

To characterise the regulatory information encoded by EpiCops, we first examined their organisation across the 833 EpiMap samples. The 720 EpiCops showed distinct tissue-specific weight profiles across the samples (**Fig. 2a**), recapitulating biological relationships among related tissues and cell types (**Supplementary Fig. 3a,b** and **Supplementary Table 1**).

**Figure 2.**
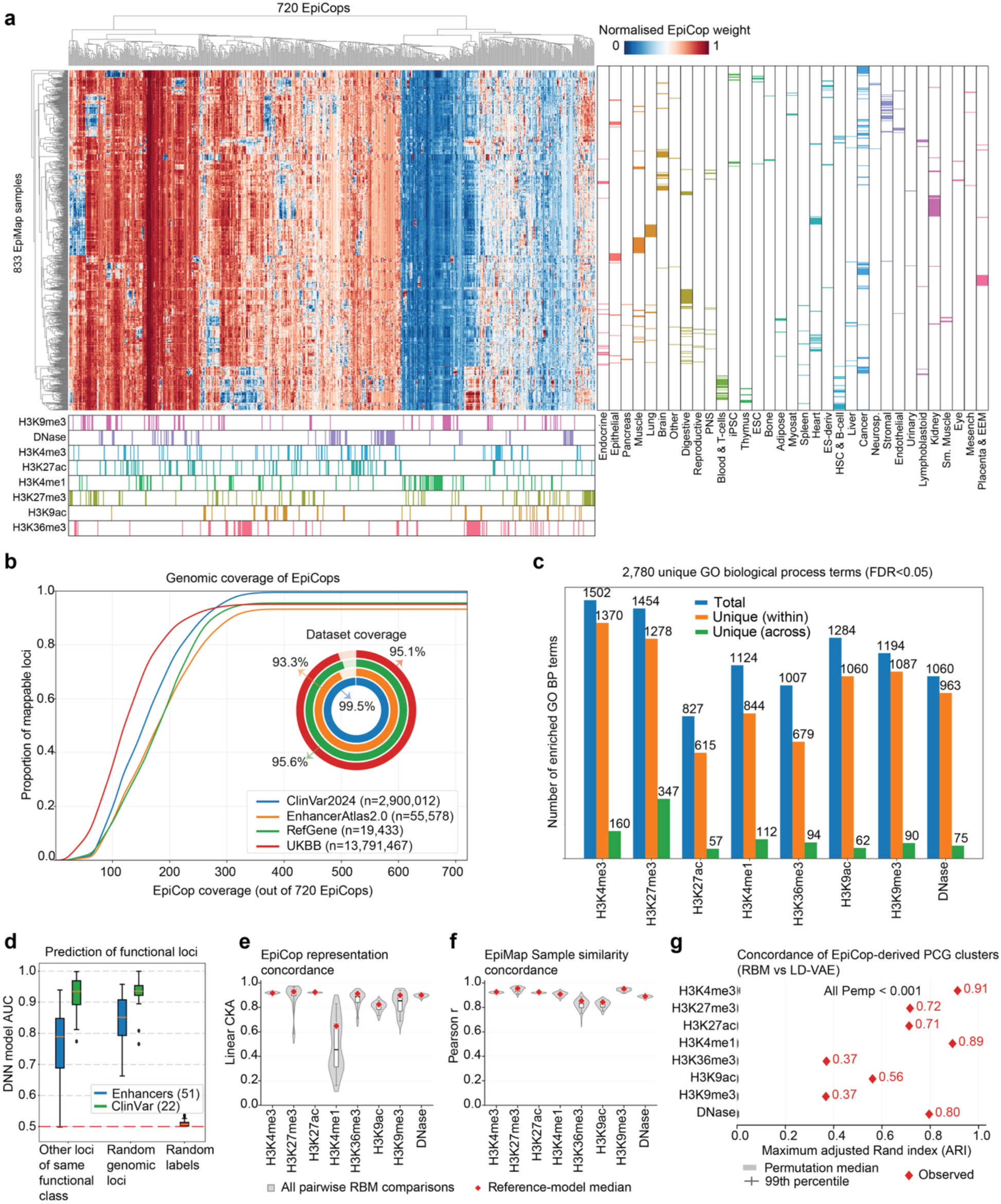
EpiCops define a coherent layer of genomic regulatory programs. **a.** EpiCop representation across 833 EpiMap samples. Heatmap shows normalised weights for 720 EpiCops. Bottom annotations indicate the chromatin mark of origin. Right annotations indicate EpiMap tissue and cell-type groups. **b.** Genomic coverage of EpiCops across loci from ClinVar 2024, EnhancerAtlas 2.0, RefGene and UK Biobank datasets. Curves show the cumulative proportion of loci (y-axis) covered as a function of the number of EpiCops that overlap the loci (x-axis). Inset shows the proportion of loci overlapping at least one EpiCop. **c.** The number of gene-ontology (GO) biological process terms enriched in protein-coding genes grouped by EpiCop combinations. GO BP enrichment was tested using Fisher’s exact test with Benjamini-Hochberg correction (FDR<0.05). Bars show all enriched terms (Total), terms unique among EpiCop combinations within each chromatin mark (Unique, within), and terms unique across chromatin marks (Unique, across). **d.** Classification of functional genomic loci using EpiCop profiles. Multi-layered neural networks were trained on loci representing tissue-specific enhancer from 51 different tissue groups, and pathogenic variants from 22 ClinVar disease groups. Performance was compared for other loci of the same function, 10,000 random genomic loci and random associations. **e.** Concordance of EpiCop representations across 20 independent RBM initialisations, quantified by linear centred kernel alignment (CKA). The plot shows all pairwise model comparisons. Red diamonds indicate the median concordance between the reference model used to generate EpiCops for the main analysis and the remaining randomly initiated models. **f.** Stability of EpiMap sample relationships across 20 independent RBM initialisations. Pairwise sample similarity was calculated from EpiCop representations, and concordance between model fits was measured by Pearson correlation of the upper triangles of the sample-by-sample similarity matrices. Red diamonds indicate the reference-model medians. **g.** Concordance of protein-coding gene (PCG) clusters derived from RBM and linear-decoded variational autoencoder (LD-VAE) representations. PCGs were clustered across Leiden clustering resolutions ranging between 0.2 and 10.0, and concordance was summarised by the maximum adjusted Rand index (ARI) across resolution pairs. Red diamonds show observed values. grey squares and crosses show the median and 99th percentile of permutation null distributions, respectively.

We next asked whether individual EpiCops represented distinct regulatory programs or redundant representations of similar chromatin activity. Pairwise cosine similarity between EpiCop weight profiles was generally low, indicating that individual EpiCops capture distinct patterns of activity across tissues and cell states (**Supplementary Fig. 3c**). To examine their genomic distributions, we converted continuous genome-wide EpiCop scores into binary occupancy using a sigmoid probability threshold of 0.5. The resulting genomic coverage varied substantially across chromatin modalities: EpiCops derived from H3K4me1, H3K9me3, H3K36me3 and H3K27me3 occupied the genome more broadly, whereas H3K27ac-, H3K9ac-and H3K4me3-derived EpiCops showed more restricted genomic distributions, reflecting the characteristic deposition patterns of these chromatin marks across the genome (**Supplementary Fig. 3d**).

Pairwise comparison of binarised genomic occupancy profiles revealed substantial co-occupancy among EpiCops derived from the same chromatin mark (**Supplementary Fig. 3e**). Co-occupancy was also evident across functionally related marks, including H3K4me1 and H3K27me3, which are associated with poised enhancers, and the active marks H3K4me3, H3K9ac and H3K27ac. The stronger overlap observed within individual marks reflects their shared genomic distribution, whereas EpiCops remain distinguished by their activity profiles across tissues and cell states. Accordingly, EpiCops represent distinct regulatory programs that can converge on the same genomic regions despite differing in their cellular activity patterns. Their genomic localisation further reflected the expected distributions of the underlying chromatin marks (**Supplementary Fig. 3f**). For example, H3K36me3-derived EpiCops were preferentially represented within exons and introns of the gene body.

EpiCops collectively provide broad coverage of functional genomic loci, with at least 93% of ClinVar pathogenic variants, EnhancerAtlas 2.0 enhancers, NCBI protein-coding genes (PCGs) and UK Biobank common variants associated with at least one EpiCop (**Fig. 2b**). Coverage increased rapidly with the number of EpiCops considered and approached saturation by approximately 300 EpiCops across most genomic datasets, indicating that individual loci are represented by specific combinations of regulatory programs rather than uniformly across all 720 EpiCops (**Fig. 2b**).

We next tested whether EpiCops were associated with distinct biological functions. For each chromatin mark, PCGs were grouped according to combinations of EpiCops present at their transcription start sites. Across these groups, we identified enrichment for 2,780 non-redundant Gene Ontology biological-process terms (**Fig. 2c**) and 174 KEGG pathway terms (**Supplementary Fig. 3g**). Most enriched GO terms were specific to individual EpiCop combinations, with each capturing a selective fraction of GO biological processes and KEGG pathways (**Supplementary Fig. 3h,i**). However, relatively fewer terms were unique to a single mark, indicating that shared biological functions are encoded through coordinated epigenetic programs across chromatin marks (**Fig. 2c**). Together, these results reveal a modular functional organisation in which individual EpiCop combinations encode specific biological functions, while related functions can be represented through different chromatin contexts.

Next, we asked whether EpiCops contained sufficient information to distinguish genomic loci with known biological functions. Using EpiCop values as input features, we trained multilayer neural network binary classifiers for 35,625 enhancers spanning 51 tissue and cell types and 2,386 pathogenic variants spanning 22 diseases. For each tissue or disease, loci belonging to the target group were distinguished either 1) from loci of the same functional class assigned to other tissues or diseases, or 2) from 10,000 randomly selected genomic loci. EpiCop-based models achieved high classification performance across both enhancer and disease-variant datasets, compared to randomised label assignments (**Fig. 2d**). These results indicate that EpiCops capture regulatory information specific to tissue, cell-type and disease contexts, providing a common representation for discriminating biologically distinct genomic loci.

We next assessed the robustness of EpiCop representations to stochastic model initialisation by retraining each chromatin mark-specific RBM model 20 times with independent random states. Concordance between the resulting sample-by-EpiCop representations was high for most chromatin marks, as quantified by pairwise centred kernel alignment (CKA) across all model fits (**Fig. 2e**). Despite greater variability in H3K4me1, relationships among EpiMap samples were strongly preserved across independently trained models for all chromatin marks, as measured by Pearson’s correlation between EpiCop-derived sample-similarity matrices (**Fig. 2f**). Together, these analyses demonstrate that EpiCop-derived biological structure is highly reproducible across independent RBM model initialisations, with sample relationships remaining stable even when the precise latent representation varies.

Finally, we asked whether EpiCop-derived regulatory structure could be recovered using an alternative representation-learning architecture. Independent epigenomic representations were learned using a linearly decoded variational autoencoder (LD-VAE), with the latent dimensionality for each chromatin mark matched to the number of EpiCops selected for the corresponding RBM. These representations were then used independently to cluster PCGs. Gene clusters derived from RBM and LD-VAE representations showed significantly greater concordance than expected from 1,000 random permutations for every chromatin mark (empirical p<0.001, **Fig. 2g**). Concordance was highest for H3K4me3 (maximum adjusted Rand index, ARI=0.91), H3K4me1 (ARI=0.89) and DNase (ARI=0.80), indicating particularly strong recovery of PCG-associated regulatory structure for these chromatin marks using independent machine learning architectures. Taken together, these results show that EpiCops capture functionally coherent regulatory structure that is reproducible across independent model fits and recoverable using a distinct representation-learning architecture.

### EpiCops recover biological structure across genomic and disease contexts

Having demonstrated that EpiCops capture reproducible regulatory programs with distinct functional associations, we next asked whether they could recover known biological structure from genomic loci with tissue or disease associations. As an initial test, we examined variants associated with atrial fibrillation (AF), a disease with a well-established cardiac regulatory component^31^. The two most enriched EpiCops among AF-associated variants showed marked heart-specific representation, with heart tissue scores substantially higher relative to the distribution across non-heart tissues (**Fig. 3a**). AF variants were significantly enriched in these two H3K4me1-derived EpiCops compared with other H3K4me1-associated genomic regions (odds ratio=1.58, p=0.003). More broadly, top 30 enriched-EpiCops were dominated by H3K4me1-derived programs, indicating a substantial contribution of enhancer-associated regulatory activity to AF. Interestingly, while the two highest-ranked EpiCops showed strong heart-specific activity, most of the remaining EpiCops were not cardiac-specific, indicating that AF-associated variants also span regulatory programs across non-cardiac tissue contexts. Together, these results show that EpiCops recover the expected cardiac regulatory architecture of AF, while also revealing extra-cardiac regulatory programs that may contribute to AF risk through systemic processes shared across tissues^32^.

**Figure 3.**
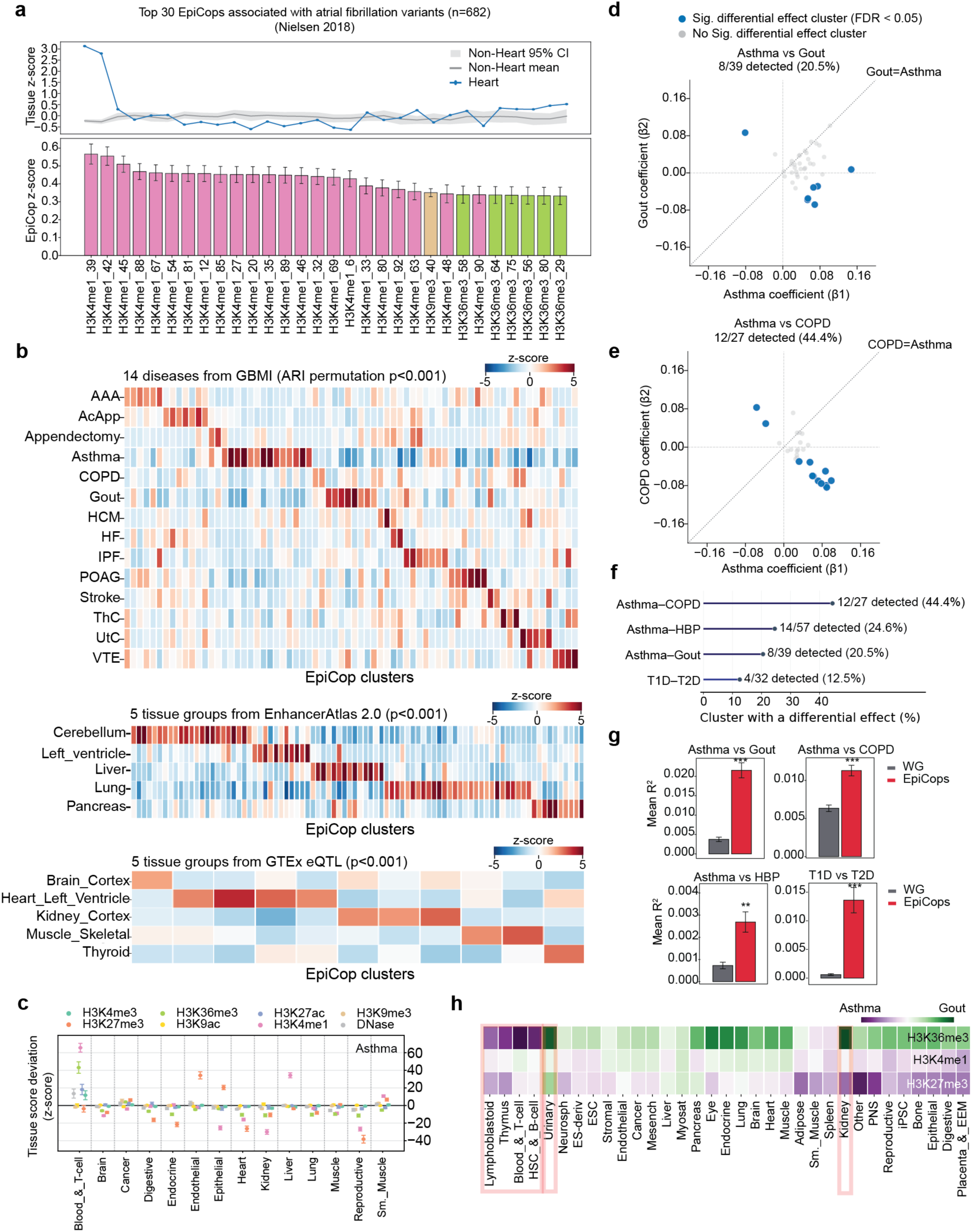
EpiCops recover biological structure across diverse genomic and disease contexts. **a.** Top 30 EpiCops enriched among 682 atrial fibrillation-associated variants. Top panel shows heart-specific tissue z-scores (blue) compared with the mean and 95% confidence interval across non-heart samples (grey). Bottom panel shows mean EpiCop enrichment z-scores observed among AF-associated variants, with the bar representing ± standard error of mean. **b.** EpiCop-based clustering of composite genomic loci from 14 GBMI diseases (top), five EnhancerAtlas 2.0 tissue-specific enhancer groups (middle) and five GTEx tissue-specific eQTL groups (bottom). Loci assigned to multiple biological contexts were excluded. Enrichment s-score and ARI permutation significance were calculated relative to 1,000 random permutations. AAA, abdominal aortic aneurysm; AcApp, acute appendicitis; COPD, chronic obstructive pulmonary disease; HCM, hypertrophic cardiomyopathy; HF, heart failure; IPF, idiopathic pulmonary fibrosis; POAG, primary open-angle glaucoma; ThC, thyroid cancer; UtC, uterine cancer; VTE, venous thromboembolism. **c.** Tissue-specific epigenetic profiles of EpiCop clusters enriched for asthma (z>2). Points show the deviation of variant-level tissue scores from the genome-wide median for each chromatin mark and tissue group. Error bars indicate 95% confidence intervals. **d-e**. Differential associations of EpiCop-partitioned variants for asthma versus gout (d) and asthma versus chronic obstructive pulmonary disease (COPD) (e). Points show test-set effect size coefficients for cluster-specific partitioned polygenic risk scores for each constituent disease. The diagonal denotes equal effects between diseases, while horizontal and vertical dashed lines indicate zero effect for the corresponding disease. Differential effects were assessed from between-disease coefficient differences using the sum of the coefficient standard errors. Benjamini-Hochberg correction across clusters was applied within each disease pair. Blue indicates clusters with FDR<0.05. **f.** Proportion of EpiCop clusters showing significant differential effects (FDR<0.05) across four disease pairs. **g.** Predictive performance of EpiCop-partitioned versus whole-genome (WG) polygenic scores across the four disease-pair comparisons. Bars show mean R^2^±standard error of mean. Asterisks indicate significant differences between models (** P<0.01, *** P<0.001). **h.** Tissue-specific regulatory profiles of EpiCops contributing to asthma-and gout-associated clusters.

We next asked whether EpiCops could recover known biological structure from a mixed set of genomic loci originating from distinct disease or tissue contexts. To this end, we constructed three independent composite datasets comprising disease-associated variants from 14 diseases^33^, tissue-specific enhancers^34^ and eQTLs^35^ from 5 distinct tissue groups. Within each dataset, loci shared across disease or tissue categories were removed before pooling, such that each retained locus was uniquely assigned to a single disease or tissue of origin. The resulting non-overlapping loci were pooled and clustered solely on the basis of their EpiCop profiles. The EpiCop-derived clusters showed significantly greater correspondence with the known disease or tissue labels than expected from 1,000 random permutations, as quantified by adjusted Rand index (ARI, empirical p<0.001 for all three datasets, **Fig. 3b**).

To characterise the tissue-level epigenetic regulation underlying this clustering structure, we next projected EpiCop profiles from disease-enriched clusters (z>2) onto their corresponding tissue and chromatin contexts. For each disease and each chromatin mark, tissue scores at disease-associated loci were expressed as deviations from the corresponding genome-wide median. Asthma-associated variants showed strong deviations across multiple chromatin programs, including H3K4me1-and H3K36me3-derived EpiCops, in blood and T-cell contexts, consistent with the prominent immune component of asthma (**Fig. 3c**). Across the broader GBMI diseases, disease-associated loci exhibited distinct combinations of tissue- and chromatin-specific deviations from the genome-wide median (**Supplementary Fig. 4**). For example, uterine cancer-associated loci showed pronounced H3K9me3-associated deviations in reproductive tissues, consistent with the established role of repressive chromatin dysregulation in endometrial malignancy^36, 37^. Idiopathic pulmonary fibrosis and venous thromboembolism both showed prominent endothelial-associated regulatory deviations, consistent with the contribution of endothelial dysfunction to both conditions^38, 39^. Together, these analyses show that EpiCop-derived clustering recovers biological structure across distinct classes of genomic loci, while providing a tissue-level epigenetic representation for the interpretation.

Having shown that EpiCops recover disease- and tissue-associated structure at the locus level, we next asked whether this information could help separate distinct genetic components within heterogeneous disease. In real-world disease cohorts, cases may comprise biologically distinct subgroups, such that subgroup-specific genetic associations are diluted when analysed together. This can limit the ability of standard genetic risk scores to distinguish between underlying disease components. We therefore constructed composite disease cohorts by pooling cases from pairs of distinct diseases and performed GWAS without using the constituent disease identities during variant discovery. Variants associated with the composite phenotype were then partitioned according to their EpiCop profiles, and cluster-specific partitioned polygenic scores (pPGSs) were evaluated in held-out individuals for their ability to distinguish the two constituent diseases. This design provided a controlled setting to test whether EpiCop-based partitioning could separate subsets of composite-associated variants carrying distinct disease-specific genetic effects.

We first tested whether EpiCop-defined variant clusters captured genetic risk for one constituent disease over the other in held-out individuals. For each cluster, we compared its association with the two diseases using effect estimates derived separately from each disease, and classified clusters with significantly different effects between the two diseases as disease divergent. In the asthma-gout composite (genetic correlation *r*_g_=0.14), 8 of 39 EpiCop-derived clusters (20.5%) showed differential effects, with several exhibiting opposing effect directions between asthma and gout (**Fig. 3d**). Furthermore, a strong separation was also observed for the pathologically related asthma-COPD pair (*r*_g_=0.66), in which 12 of 27 clusters (44.4%) showed significantly differential disease effects (**Fig. 3e**). Differential clusters were also detected for asthma-high blood pressure (14 of 57 clusters or 24.6%, *r*_g_=0.162), and for type 1 and type 2 diabetes (4 of 32 clusters or 12.5%, *r*_g_=0.002), demonstrating that EpiCop-defined clusters can identify subsets of variants with divergent disease-specific effects across composite diseases spanning a wide range of genetic correlation (**Fig. 3f**). Across all four disease pairs, jointly fitting EpiCop-defined pPGS components explained significantly more variation in distinguishing the constituent disease subtypes than an unpartitioned PGS constructed from the same variants (**Fig. 3g**). Beyond this improvement in predictive resolution, the resulting clusters also captured distinct tissue-level epigenetic associations. For example, in the asthma-gout composite, asthma-associated clusters preferentially captured immune and haematopoietic programs, whereas gout-associated clusters showed stronger kidney-and urinary-associated signals, consistent with the known tissue-level biology of these diseases (**Fig. 3h**). Together, these analyses provide a proof of concept that EpiCop-based clustering can resolve distinct genetic mechanisms within a heterogeneous disease by partitioning associated variants according to their underlying tissue-level epigenetic programs, improving discrimination of biologically distinct disease components compared with unpartitioned models.

### EpiCops reveal heterogeneous regulatory architectures in type 2 diabetes

We next asked whether the same EpiCop representation could reveal biological heterogeneity within a single complex disease, where the underlying mechanisms are not known. We focused on type 2 diabetes (T2D), a heterogeneous disorder involving metabolic and vascular dysfunction across multiple tissues and organ systems. Previous genetic subtyping studies have partitioned T2D-associated variants according to their associations with auxiliary cardiometabolic traits, identifying components linked to distinct disease mechanisms including insulin deficiency, obesity, lipodystrophy and lipid metabolism^15, 16^. Here, we used an unsupervised approach to cluster T2D-associated variants reported by Smith et al.^15^ using only their EpiCop profiles, without incorporating phenotypic information into cluster definition. Cardiometabolic traits were subsequently evaluated across the resulting clusters as independent phenotypic readouts of the biological structure recovered from epigenomic regulation.

Across 36 cardiometabolic traits, the aggregate T2D variant set showed the expected pattern of broadly positive associations with glycaemic, adiposity, lipid and cardiovascular phenotypes. EpiCop-derived clustering decomposed this aggregate signal into markedly heterogeneous profiles, with individual clusters showing selective enrichments and, in some cases, opposing effects across related phenotypes (**Fig. 4a**). These divergent association patterns reveal phenotypically distinct components of T2D genetic risk that were not apparent when variants were analysed as a single combined set.

**Figure 4.**
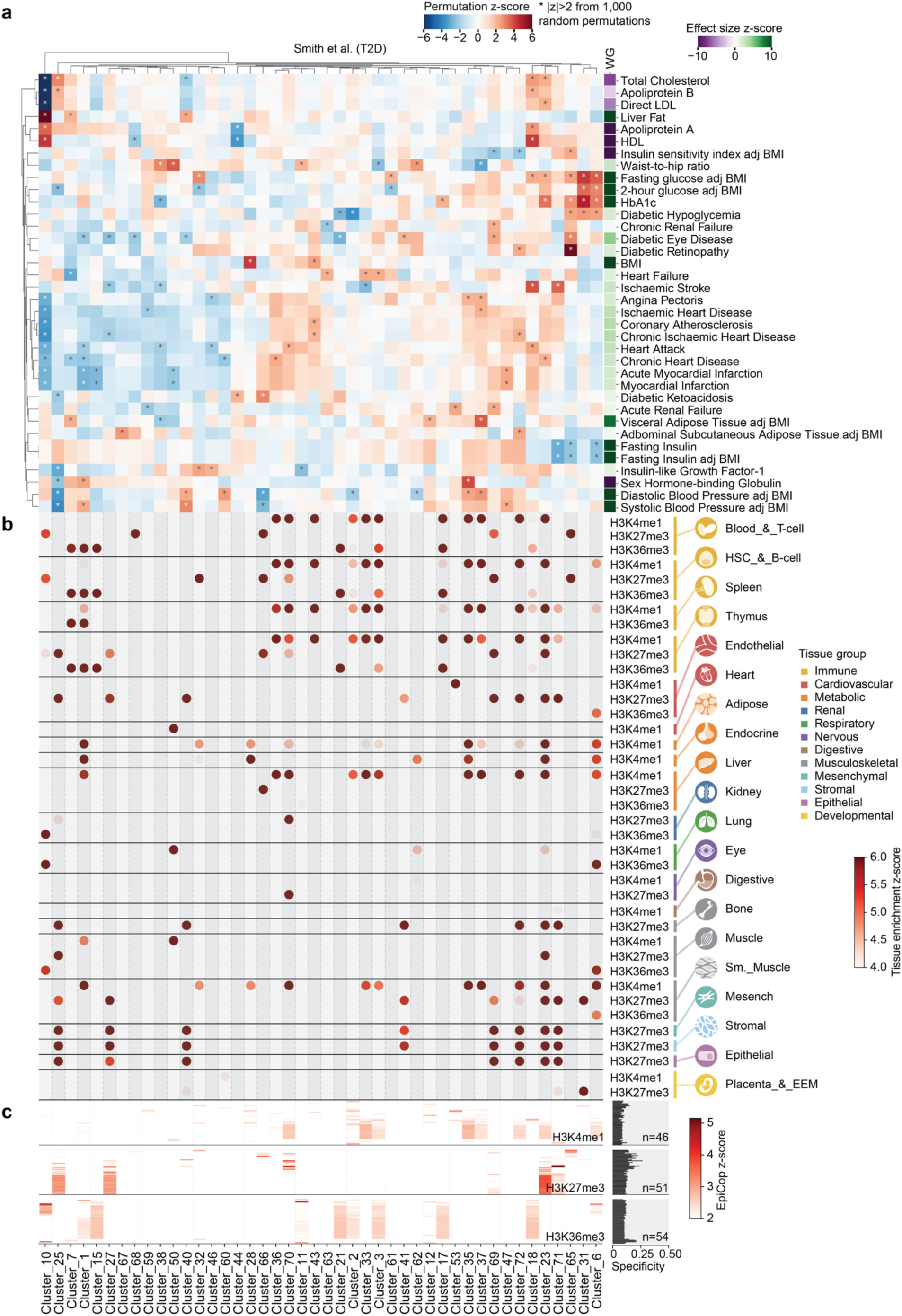
EpiCops reveal distinct sub-disease structures of type 2 diabetes. **a.** Enrichment of EpiCop-defined clusters of type 2 diabetes-associated variants across 36 cardiometabolic traits. Cluster-level enrichment z-scores were calculated from aggregated variant effect sizes relative to 1,000 random permutations. Only clusters with an absolute enrichment z-score above 2 for at least one cardiometabolic trait are shown. **b.** Tissue-specific epigenetic association with each EpiCop-defined variant cluster. Only positive enrichments with z-score above 4 are shown. **c.** Enrichment of individual EpiCops across T2D variant clusters. Only EpiCops with enrichment above z-score of 2 are shown. Bars at right indicate EpiCop cluster specificity, quantified as 1-normalised entropy, where 0 indicates broad sharing across all clusters and 1 indicates maximal cluster specificity.

We next examined the regulatory contexts underlying this phenotypic heterogeneity. Mapping the EpiCop profiles of each variant cluster onto tissue-specific chromatin programs revealed distinct combinations of tissue and chromatin activity (**Fig. 4b**). Notably, clusters associated with increased risk of angina pectoris, ischaemic heart disease and myocardial infarction showed prominent H3K4me1 enrichment across blood and T cells, haematopoietic stem and B cells, spleen and thymus. This pattern is consistent with previous evidence implicating immune-associated enhancer activity and H3K4me1-marked regulatory regions in coronary artery disease and atherosclerotic inflammation^40^. By contrast, clusters showing reduced cardiovascular effects, including cluster 10, showed little corresponding H3K4me1 enrichment but prominent H3K27me3 signals across the same tissue groups. Reduced H3K27me3 has previously been observed in inflammatory monocytes and advanced atherosclerotic lesions^9, 10, 40^, supporting a link between altered repressive chromatin states and cardiovascular disease.

More broadly, clusters mapped to different combinations of metabolic, vascular, cardiac, renal and immune tissue contexts, indicating their distinct cardiometabolic trait associations do not reflect a single tissue of action, but instead correspond to distinct combinations of epigenomic regulation across multiple tissues and organ systems.

This organisation was also evident at the level of individual regulatory programs. Enriched EpiCops differed in how selectively they were represented across T2D variant clusters, as quantified by Shannon entropy-based cluster specificity scores (**Fig. 4c**). Some EpiCops recurred across several clusters, whereas others were restricted to specific subsets, indicating that T2D-associated variants are organised through partially shared yet distinct combinations of regulatory programs. These patterns further support a modular regulatory basis underlying the phenotypic heterogeneity of T2D.

We next tested whether the T2D sub-disease structure revealed by EpiCop was robust to alternative clustering procedures and latent feature-extraction architectures. Among alternative clustering algorithms matched to the same number of clusters, Leiden community detection showed substantially greater concordance with the EpiCop-derived clustering than either k-means or hierarchical clustering (ARI=0.809, versus 0.248 and 0.224 respectively, **Supplementary Fig. 5a**), indicating that the T2D sub-disease structure is largely preserved by graph-based community clustering^30, 41^.

Using the previously generated LD-VAE representations, with the number of latent features matched to the corresponding RBM-derived EpiCops for each chromatin mark, we next repeated the T2D variant clustering analysis within the same Leiden clustering framework. At equivalent clustering resolutions, LD-VAE representations produced a larger number of clusters, whereas RBM-derived EpiCops achieved higher silhouette scores across most cluster numbers, indicating a more compact and clearly separated partition of T2D-associated variants (**Supplementary Fig. 5b,c**). Despite these differences in cluster structure, both representations recovered heterogeneous cardiometabolic trait associations with broadly comparable effect-size ranges (**Supplementary Fig. 5d**). Together with the cross-architecture concordance observed at protein-coding gene loci, these findings show that the regulatory heterogeneity resolved within T2D is not specific to the RBM feature-extraction architecture, but can also be recovered from independently learned LD-VAE representations, supporting the biological robustness of the T2D sub-disease structure revealed by EpiCops.

### Disease-agnostic EpiCops benchmark favourably against disease-informed variant partitioning approaches

To benchmark EpiCop-based variant partitioning, we compared its performance with existing approaches that incorporate disease-relevant phenotypic or tissue information. We compared EpiCops with five comparator frameworks spanning auxiliary cardiometabolic trait-based clustering^15^, tissue-specific open chromatin^20^, joint pleiotropic and epigenomic modelling^42^, phenotype-derived latent components^43^ and disease-specific epigenomic partitioning^44^, evaluated in T2D and asthma (**Fig. 5a**). By contrast, EpiCops are learned independently of disease context from consortium-scale epigenomic data and subsequently applied to disease-associated variants without phenotypic or disease-specific priors. For each head-to-head comparison, analyses were performed on the same variant set available to the corresponding comparator, and cluster-level enrichment for disease-relevant traits (**Supplementary Table 2**) was quantified relative to null distributions generated from 1,000 random permutations.

**Figure 5.**
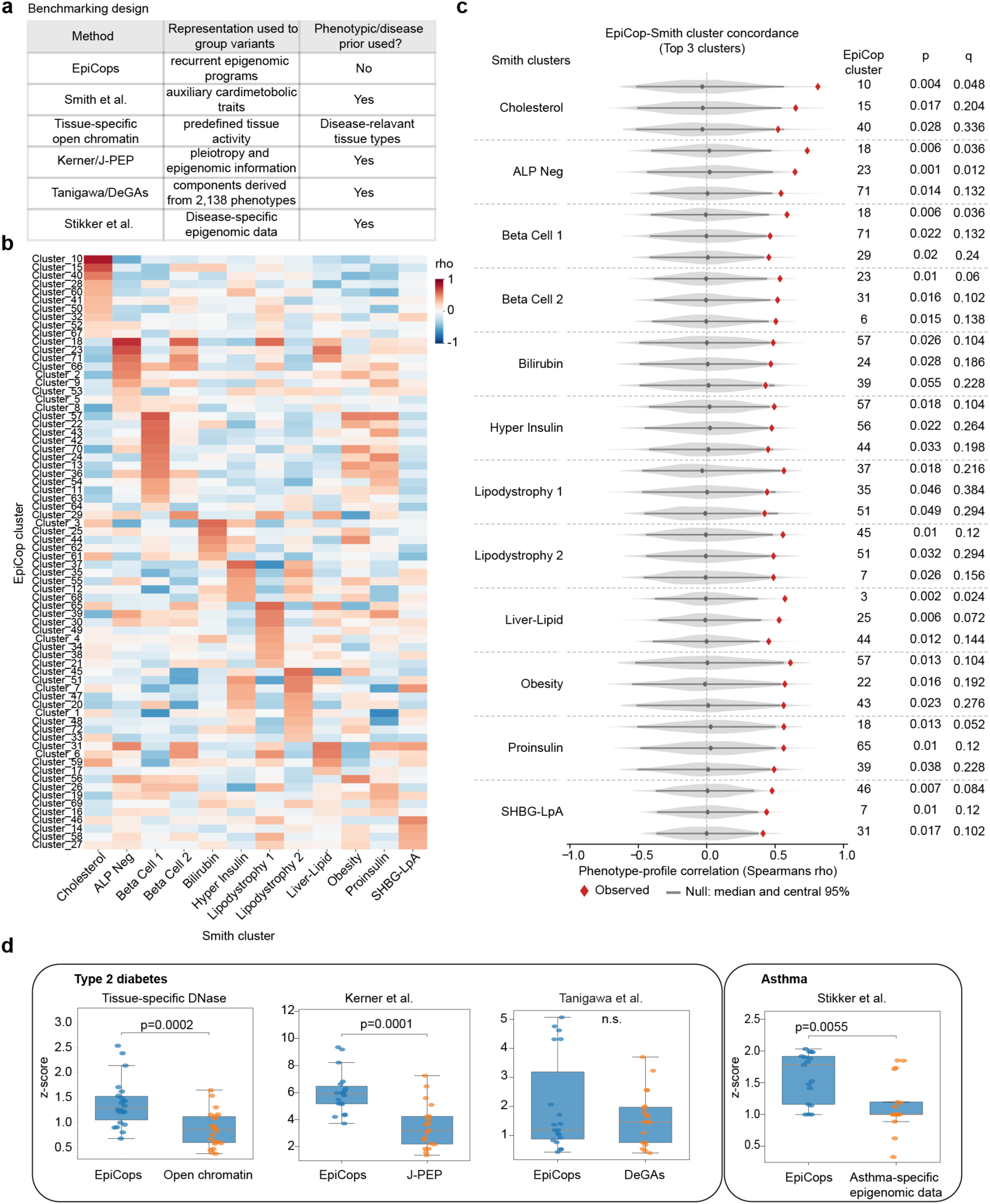
Benchmarking of EpiCop-derived T2D variant clusters. **a.** Benchmarking approaches and the information used to group variants. **b.** Spearman correlation coefficients (rho) between phenotype associations of 72 EpiCop clusters and 12 Smith clusters. **c.** Concordance of EpiCop-derived clusters with T2D cardiometabolic phenotype-based clusters (Smith et al.). Top three EpiCop matches for each Smith cluster component, ranked by Spearman correlation across the 36 cardiometabolic traits. Red diamonds indicate observed correlations. Grey violins, dots and bars show null distributions, medians and central 95% intervals, generated from 1,000 cluster label permutations. One-sided empirical p-values and Benjamini-Hochberg-adjusted q-values are shown. **d.** Maximum cluster-level phenotype enrichment z-scores for EpiCops and comparator methods in T2D and asthma. Each point represents a single disease-relevant phenotype. Signifance was assessed with Wilcoxon ranksum test.

We first tested whether EpiCop-based clustering could independently recover established components of T2D heterogeneity identified by phenotype-informed approaches. Smith et al. defined 12 T2D genetic components from auxiliary cardiometabolic trait associations, providing a direct benchmark against the 72 EpiCop-derived clusters. Comparison of their cardiometabolic association profiles revealed selective concordance between individual EpiCop-derived and Smith clusters (**Fig. 5b**). Among the strongest matches, observed phenotypic correlations generally exceeded permutation-derived expectations, with most showing nominal empirical significance and a subset remaining significant after multiple-testing correction (**Fig. 5c**). Thus, despite using no cardiometabolic phenotypes for cluster definition, EpiCop-based partitioning recovered variant groups corresponding to previously identified phenotype-defined components of T2D heterogeneity.

We then benchmarked whether EpiCop-partitioned variant clusters are enriched for disease-relevant phenotypes. For each trait, we compared the strongest cluster-level enrichment obtained with EpiCops against that obtained with the corresponding comparator approach, both quantified as the maximum permutation-standardised z-score (**Fig. 5d**). In T2D, EpiCop-derived clusters showed significantly greater enrichment across cardiometabolic traits than clusters defined by tissue-specific open chromatin (p=0.0002) or the joint pleiotropic and epigenomic modelling (p=0.0001), while performing comparably to latent genetic components derived from GWAS associations across 2,138 UK Biobank phenotypes. In asthma, EpiCop-derived clusters showed greater enrichment across asthma-relevant phenotypes than clusters defined using asthma-specific epigenomic annotations (p=0.0055).

Together, these benchmarking analyses show that EpiCops can recover previously established disease heterogeneity and identify phenotypically informative variant subsets without requiring prior phenotypic or disease-specific information for cluster definition. As an unsupervised, disease-agnostic regulatory representation, EpiCops performed comparably to or more strongly than approaches that explicitly incorporate disease-relevant phenotypic information, tissue-specific annotations or disease-specific epigenomic features.

### Opposing regulatory architectures define divergent cardiovascular outcomes in type 2 diabetes

To determine whether the regulatory composition of inherited T2D risk captures clinically and phenotypically relevant variation, we quantified each EpiCop-derived variant cluster as a pPGS. Jointly modelling these pPGS components explained more phenotypic variance across diverse T2D-relevant phenotypes relative to an aggregated T2D PGS constructed from the same set of variants (**Supplementary Fig. 6a**), indicating that partitioning genetic risk by regulatory programs captures information that is obscured when risk variants are aggregated into a single score.

This distinction was particularly evident for clusters 10 and 36, which showed opposing cardiometabolic profiles. Higher cluster 10 pPGS, which reflects greater weighted genetic burden from variants assigned to cluster 10, was associated with lower LDL and total cholesterol, lower triglycerides, higher HDL cholesterol, and reduced myocardial infarction and chronic ischaemic heart disease risk, whereas cluster 36 showed the opposite cardiovascular profile (**Fig. 6a** and **Supplementary Fig. 6b**). Interestingly, cluster 10 was simultaneously associated with increased liver fat and fatty liver disease, revealing opposing hepatic and cardiovascular consequences within the same component of T2D genetic risk. Together, EpiCop-based pPGS can partition an individual’s genetic risk into components governed by distinct regulatory signatures with different phenotypic effects.

**Figure 6.**
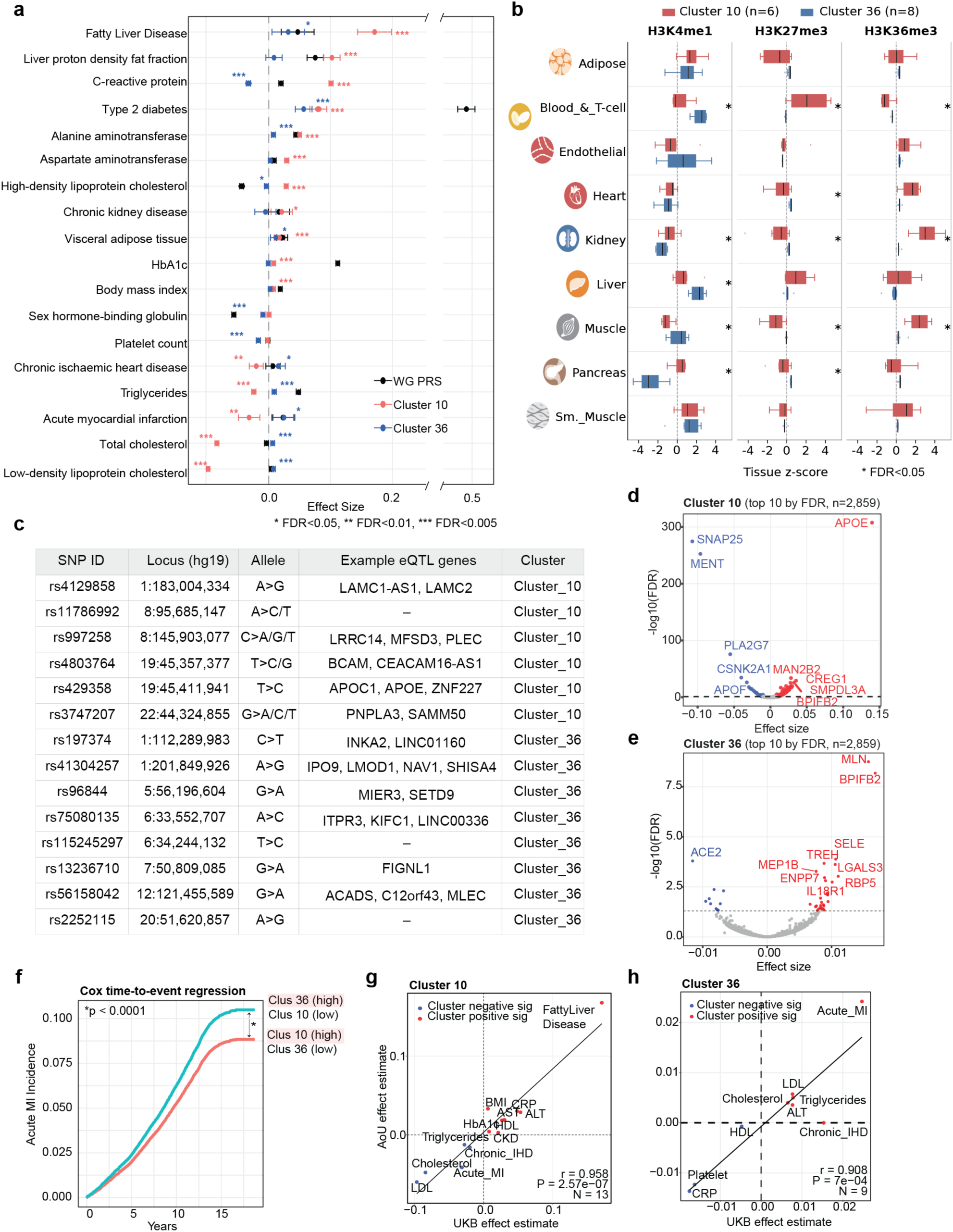
EpiCop-derived clustering identifies T2D genetic sub-structures with opposing cardiovascular outcomes. **a.** Comparison of effect sizes for selected cardiometabolic traits between clusters 10 and 36 using cluster-specific partitioned polygenic models. Asterisk marks indicate statistical significance compared with the whole-genome PGS (* FDR<0.05, ** FDR<0.01, *** FDR<0.005). Whole genome (WG) represents effect size for all T2D variants. **b.** Tissue-specific epigenetic profiles of the cardioprotective (Cluster 10) and high cardiovascular-risk (Cluster 36) clusters. Only chromatin marks represented by EpiCops with noticeable variability among T2D-associated variants (median absolute deviation>0.5) are shown. Values represent mean tissue-level EpiCop z-scores, with zero indicating the genome-wide mean. Asterisks indicate significant differences between these two clusters (Wilcoxon rank-sum test, FDR<0.05). **c.** Variants assigned to clusters 10 and 36, including genomic coordinates, alleles (ref>alt) and representative eQTL-linked genes. **d-e.** Association of top 10 protein biomarkers associated with clusters 10 (d) and 36 (e). **f**. Cox time-to-event regression analysis for year-to-acute myocardial infarction incidence in UK Biobank accounting for overall T2D risk and diagnosis status. **g-h**. Concordance of cluster-specific cardiometabolic trait effects between UK Biobank and All of Us for clusters 10 (g) and 36 (h), measured by Pearson’s correlation coefficient. Each point represents a single trait. Pearson correlation coefficients are shown.

We further examined the regulatory and molecular profiles underlying these divergent phenotypic effects. Cluster 10 showed greater H3K27me3-associated representation in blood and T-cell contexts and greater H3K36me3-associated representation in kidney and muscle, whereas cluster 36 showed greater H3K4me1-associated representation across immune and hepatic contexts (**Fig. 6b**). The clusters also differed in their constituent loci. Cluster 10 contained variants with eQTL links to PNPLA3 and APOE^35^, whereas cluster 36 included variants linked to genes such as LMOD1, MIER3 and ACADS (**Fig. 6c**). The two clusters were further distinguished by their plasma-protein associations^45^, with cluster 10 linked to proteins involved in lipid transport and metabolism, including SMPDL3A and APOE^46–48^, whereas cluster 36 was associated with proteins implicated in vascular and inflammatory dysfunction, including SELE and LGALS3^49^ (**Fig. 6d,e**). This lipid-related molecular profile provides a potential mechanistic context for the opposing hepatic and cardiovascular effects observed for cluster 10. This is consistent with studies demonstrating that increased hepatic lipid retention can reduce the export of ApoB-containing lipoproteins, lowering circulating atherogenic lipids and coronary risk^50^. Together, these findings indicate that the opposing cardiometabolic effects of clusters 10 and 36 are associated with distinct combinations of genetic loci, tissue and chromatin context, and circulating molecular profiles, linking distinct regulatory profiles to divergent physiological manifestations of T2D genetic risk.

We next tested whether these divergent cardiometabolic risks translated into prospective clinical outcomes. UK Biobank participants with high cluster 10 and low cluster 36 pPGS (cluster 10 pPGS ≥ P70 and cluster 36 pPGS ≤ P30) had a 17% lower hazard of incident myocardial infarction than participants with the reciprocal genetic profile (Cox proportional hazards model, p<0.0001, **Fig. 6f**). Consistent with the opposing associations with fatty liver traits, individuals stratified by these opposing risk clusters (clusters 10 and 25) showed significantly different cumulative incidence of non-alcoholic fatty liver disease (NAFLD) during follow-up (log-rank p<0.0001, Cox HR=1.39, **Supplementary Fig. 6c**). Notably, cluster 10 was associated with increased hepatic risk despite its previously observed protective cardiovascular profile. Together, these results genetically dissociate hepatic and cardiovascular complications within T2D susceptibility, showing that a regulatory component contributing to T2D risk can have divergent consequences across organ systems^49^.

We then assessed whether these cluster-specific effects replicate beyond UK Biobank using the independent All of Us cohort^51^. Across significant cardiometabolic associations, cluster-specific pPGS effect estimates showed strong cross-cohort concordance for both cluster 10 (*r*=0.958, p=2.57*10⁻⁷, *N*=13 traits, **Fig. 6g**) and cluster 36 (*r*=0.908, p=7*10⁻⁴, *N*=9 traits, **Fig. 6h**). Similar concordance across the broader set of continuous and binary traits was also observed (**Supplementary Fig. 6d,e)**. Together, these findings link distinct cellular and epigenomic programs underlying T2D-associated variation to molecular and phenotypic profiles and prospective clinical outcomes, providing a mechanistic basis for clinically meaningful heterogeneity within a shared disease diagnosis.

Finally, we asked whether phenotypically distinct regulatory components could also be identified beyond T2D. EpiCop-based partitioning of asthma-associated variants revealed clusters with distinct associations across age-of-onset, respiratory, metabolic and circulating immune-cell phenotypes (**Supplementary Fig. 7a,b**). Consistent with T2D, incorporating EpiCop-derived asthma clusters into partitioned polygenic models increased phenotypic variance explained relative to the corresponding aggregate model (**Supplementary Fig. 7c**). These results indicate that phenotypically distinct regulatory components identified by EpiCops extend across diverse, independent complex diseases.

## Discussion

Our findings identify a layer of genomic organisation in which genomic loci are related through recurrent patterns of chromatin activity across tissues and cell states. Genomic variation is commonly interpreted through nearby genes, pathways, individual regulatory elements or enrichment within predefined tissues and cell types^14, 42, 52^. EpiCops provide a complementary representation in which loci are grouped according to regulatory patterns that recur across diverse cellular contexts. These programs therefore capture relationships between genomic regions that are not apparent from individual epigenomic profiles or annotations, providing a common regulatory coordinate system through which genomic loci can be interpreted according to the cellular and chromatin contexts in which they operate.

Importantly, these regulatory programs are neither equivalent to cell identity nor restricted to individual tissues. Instead, cell identity and function arise from combinations of programs that can recur across tissues and cellular states, whereas loci within the same tissue can belong to different programs depending on their chromatin context. This distinction was particularly evident in T2D, where multiple variant clusters involved immune-associated regulatory activity but differed in their underlying chromatin programs. Individual groups also spanned across immune, metabolic, vascular and other tissues. EpiCops therefore provide an axis of biological organisation complementary to tissue or cell-type annotation, capturing both the cellular context where genomic loci repeatedly act and how they are regulated in coordination. Such a representation may be particularly useful for interpreting complex biological processes that emerge through interactions across multiple cell states and organ systems.

Complex disease provides a stringent setting in which to test whether these regulatory programs capture biologically meaningful information. Existing approaches to disease stratification commonly use clinical or molecular phenotypes measured after disease developed and are therefore subject to reverse causation^53, 54^. While genetic subtyping approaches can resolve more upstream mechanisms underlying disease heterogeneity using predefined pathways, functional annotations or auxiliary trait associations^15, 16^, they rely heavily on the biological information used to define them. EpiCops were instead learned independently of disease and subsequently used to group disease-associated variants according to their underlying regulatory context. In composite phenotypes with known constituent diseases, this organisation separated mixed genetic signals according to their ground-truth biological origins using regulatory information alone. Thus, aggregate genetic models can conceal mechanistically coherent relationships between disease-associated variants that become apparent when genetic variation is represented within latent regulatory space.

T2D demonstrated that this regulatory organisation extends to biological heterogeneity within a single complex disease. Despite being derived without phenotypic information, EpiCops recovered established components of T2D heterogeneity identified by phenotype-informed approaches, while revealing additional differences in tissues and chromatin states. Variants contributing to the same disease were partitioned into distinct regulatory profiles associated with divergent, and in some cases opposing cardiometabolic effects. At the individual level, these regulatory components generated distinct partitioned genetic risk profiles whose phenotypic associations were reproduced across independent populations. Particularly notable was an architecture represented by cluster 10, in which increased susceptibility to hepatic steatosis occurred alongside favourable lipid profiles and reduced cardiovascular risk. This finding provides a regulatory context for the established genetic dissociation between hepatic steatosis and coronary disease, where genetic mechanisms that increase liver fat do not necessarily confer the cardiovascular consequences expected from their metabolic phenotype. Rather than defining uniformly favourable or adverse disease subtypes, regulatory decomposition therefore separates components of T2D liability whose effects can diverge across tissues, organ systems and clinical outcomes.

These observations have implications for how inherited risk is represented. Conventional aggregate genetic models quantify the cumulative contribution of disease-associated alleles into a single measure of liability, even if they act through different biological mechanisms. Our findings suggest that genetic risk is characterised not only by its magnitude but also by its regulatory composition, as variants contributing to the same clinical phenotype can be distributed across distinct regulatory architectures with different cellular, molecular and physiological consequences. EpiCop decomposition preserves information about this composition, providing a potential basis for understanding how shared susceptibility to a complex disease can give rise to heterogeneous manifestations and subsequent disease trajectories. This framework remains compatible with stratagenic models of complex disease^8^, but suggests that distributed genetic effects need not be biologically unstructured. Instead, a large number of individually modest genetic effects may converge through a more restricted set of recurrent regulatory contexts that connect inherited variation to cellular function.

Several features of the current framework define both its scope and its limitations. EpiCops represent recurrent regulatory relationships learned from existing epigenomic reference datasets and therefore depend on the diversity of tissues, cellular states and chromatin modalities represented within these resources. Rare, transient or disease-specific regulatory states that are absent from these atlases may consequently be incompletely represented. Moreover, EpiCop assignment identifies regulatory contexts associated with genomic loci but does not itself establish the causal molecular mechanism through which individual variants influence cellular phenotype. Increasingly comprehensive epigenomic atlases, particularly single-cell and perturbational datasets, should enable these architectures to be resolved at greater cellular and functional resolution and determine whether regulatory programs inferred from population-scale reference maps correspond to regulatory states directly observed in disease-relevant cells. Together, our findings establish latent regulatory programs as a generalisable framework for organising genomic variation across cellular contexts, revealing biological structure that links inherited genetic risk to heterogeneous molecular, physiological and clinical outcomes.

## Data and code availability

Upon publication, the source code and processed data required to reproduce the analyses will be made publicly available through a permanent repository.

## Author contributions

W.J.S. conceptualised the study, developed the EpiCop methodology, performed formal analyses, and wrote the original manuscript draft. S.C.B. performed validation analyses to demonstrate application of EpiCops in complex disease across multiple population cohorts. C.S.Y.C. performed pre-processing of EpiMap consortium data. S.S. (Sophie Shen) contributed to data visualisation by generating figure schematics. Z.R. performed comparative analysis using LD-VAE. Q.Z. performed pre-processing of EpiMap consortium data. J.Z. and S.S. (Sonia Shah) provided methodological advice on variant clustering approaches and performed analysis on statistical genetic models. D.M. performed analysis on composite disease traits and provided supervision of S.C.B. M.B. provided supervision of W.J.S. N.J.P. provided overall supervision of the project and acquired funding. All authors contributed to manuscript review and editing.

## Acknowledgements

This work was supported by the National Heart Foundation of Australia (grant 106721 to N.J.P.), the Medical Research Future Fund (MRF2055599, APP2053737 and APP2016033 to N.J.P.), and the National Health and Medical Research Council of Australia (grant 2043111 to J.Z.). S.S. was supported by a Heart Foundation Future Leader Fellowship. This research has been conducted using the UK Biobank Resource under Application Number 899824.

## Methods

### Datasets

ChIP-Sequencing data from Epimap consortium were used to extract patterns of multiple epigenetic marks in the human genome^20^. These include DNase I hypersensitivity and 7 different histone tail modifications including H3K4me3, H3K27me3, H3K4me1, H3K27ac, H3K9ac, H3K9me3 and H3K36me3. All available bigWig files were used, including partially imputed epigenetic signals from diverse cellular contexts (https://epigenome.wustl.edu/epimap/data/imputed/). We used 833 QC-passed samples only as demonstrated previously^20^.

These bigwig data were converted to bedgraph format using bigWigToBedGraph program from UCSC executable download page (http://hgdownload.cse.ucsc.edu/admin/exe/), with the default parameters. Narrow peaks for H3K4me3, H3K27ac, H3K9ac, and DNase hypersensitivity were called using MACS3’s bdgpeakcall function, whereas broad peaks for H3K4me1, H3K27me3, H3K9me3, and H3K36me3 were called using bdgbroadcall function^55^. This differential peak-calling strategy ensures capturing of the distinct genomic distributions of epigenetic marks, as previously described^56^.

We binarized the epigenomic data for several advantages as previously demonstrated^21, 57, 58^; 1. increased robustness to outliers, 2. improved comparability between samples across different epigenetic marks, and 3. substantial reduction in computational load during model training, with minimal information loss. This is particularly important given the scale of the epigenomic datasets derived from the human genome.

To assess sample specificity, we counted the number of samples in which peaks for a given epigenetic mark were observed within 1 kb genomic windows. We then calculated the proportion of samples exhibiting peaks relative to the total number of samples. Similarly, we computed the proportion of peaks within each window relative to the total number of peaks for that epigenetic mark. These proportions were interpolated into a single plot to compare sample specificity across epigenetic marks at 1 kb resolution. We evaluated enrichment of peaks at FANTOM5 robust enhancers, long non-coding RNAs^27^ transcription start sites (TSSs) of RefSeq protein-coding genes (PCGs) by calculating fold enrichment relative to 20,000 randomly selected genomic peaks. Fold enrichment was computed independently for each sample and each epigenetic mark.

### Identifying epigenomic patterns using Restricted Boltzmann Machine

To extract EpiCops, we trained Restricted Boltzmann Machine (RBM) models independently for each of the 8 epigenetic marks using BernoulliRBM module implemented in Python’s scikit-learn package^59^. RBMs are shallow generative neural networks that learn latent structure through auto-associative modelling between visible and hidden layers, enabling coordinated chromatin patterns to be represented through bi-directional weights that remain directly interpretable across tissue and cell-type contexts^29^. Input data for each model were represented as a binary matrix *M* ∈ {0,1}^∼2.72*M*∗^ *^y^*, where each row corresponds to a non-overlapping 1 kb genomic window (i.e. ∼2.72 million windows in the hg19 genome where ChIP-seq peaks were observed from any samples) and each column represents one EpiMap sample. A value of 1 indicates the presence (i.e. at least 1bp overlap) of the epigenetic mark in a given genomic window for an epigenetic mark.

For each epigenetic mark, we trained models using heuristically selected hyperparameters. For each epigenetic mark, we performed grid-search optimisation over key hyperparameters, including the learning rate, random initialisation state, number of training iterations, and number of hidden nodes. We used python sklearn’s BernoulliRBM for the training.

We used the selected model settings to identify the optimal number of hidden nodes (n_components). For this step, model performance was evaluated using an 80:20 training:test split data within a five-fold cross-validation framework. For each trained RBM model, the held-out test set was reconstructed from its learned hidden representation. Specifically, the hidden representation of the test data was first obtained using rbm.transform (test_data). The reconstructed visible-layer values were then estimated by taking the dot product between this hidden representation and the learned RBM weight matrix (rbm.components_). A sigmoid transformation was applied to convert the reconstructed values into predicted probabilities, which were then binarised to generate the predicted output. Model reconstruction error was quantified as the mean squared error between the reconstructed and observed test data. Models were trained and evaluated across n_components=5-100, and the model with the lowest reconstruction error was selected to define the optimal number of hidden nodes for each epigenetic mark. In this framework, each hidden node represents a single latent epigenomic program encoded by weighted associations across biological samples. This optimisation procedure produced eight independently trained RBM models, each capturing recurrent chromatin programs specific to one epigenetic mark.

### Sensitivity and cross-architecture analyses of EpiCop representations

To assess the robustness of EpiCop representations to stochastic model initialisation, each chromatin mark-specific RBM was trained 20 times using different random states while holding the optimised number of hidden nodes, learning rate, number of training iterations and batch size constant. Each model was trained on the complete binary genomic-window-by-sample matrix, and the resulting hidden-unit weight matrix was extracted. Each weight matrix was transposed to obtain an EpiMap sample-by-EpiCop representation. Concordance between representations from independently initialised RBMs was quantified using centred kernel alignment (CKA) across all 190 unique pairwise comparisons among the 20 model fits. For visualisation, the distribution of pairwise CKA values was shown for each chromatin mark together with the median concordance between the reference model used in the primary analysis and the 19 alternative initialisations.

To independently assess preservation of relationships among EpiMap samples, a sample-by-sample Pearson correlation matrix was calculated from the EpiCop representation of each RBM model. For each pair of independently initialised models, concordance was quantified by calculating Pearson correlation between the corresponding upper-triangular off-diagonal elements of the two sample-similarity matrices. Spearman correlation was additionally calculated as a rank-based measure of concordance. As above, comparisons were performed across all 190 unique pairs of RBM fits, with the median concordance between the reference model and the 19 alternative initialisations reported. This analysis therefore assessed conservation of EpiMap sample relationships without requiring one-to-one correspondence between individual hidden units.

To assess whether biological structure derived from EpiCops was preserved across different latent-variable architectures, we compared RBM-derived representations with latent representations generated using a linearly-decoded variational autoencoder (LD-VAE). For each chromatin mark, the LD-VAE latent dimensionality was fixed to the number of EpiCops selected for the corresponding RBM, providing a matched-capacity comparison between model architectures. Protein-coding gene transcription-start-site loci were clustered independently from the RBM-and LD-VAE-derived representations using Leiden clustering across resolutions from 0.2 to 10.0 in increments of 0.2. After aligning the same genomic loci between models, adjusted Rand indices (ARI) were calculated for all pairwise combinations of RBM and LD-VAE clustering resolutions, and the maximum observed ARI was retained for each chromatin mark.

To account for selection of the resolution pair that maximised concordance, statistical significance was assessed using 1,000 permutations. In each permutation, LD-VAE locus identities were randomly permuted using a single common permutation across all resolution-specific cluster assignments, thereby preserving cluster sizes and the dependence among LD-VAE partitions across resolutions. ARI was then recalculated across the complete set of RBM–LD-VAE resolution combinations and the maximum ARI retained, generating a null distribution of permutation-calibrated maximum ARIs. Empirical p-values were calculated as

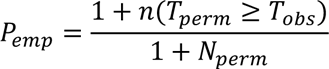

where *T_obs_* is the observed maximum ARI and *T_perm_* is the maximum ARI obtained in each permutation. The median and 99th percentile of the permutation-derived maximum ARI distribution were used to summarize the null expectation.

### Genome-wide distribution of EpiCops

We performed matrix multiplication to identify genomic regions that are associated with EpiCops. For a given epigenetic mark, let W ∈ *R^f^*^∗^ *^y^* be a weight matrix where each row represents an EpiCop and each column represents an EpiMap sample, with *f* and *y* representing the total numbers of EpiCops and EpiMap samples for that epigenetic mark, respectively. We first converted the binary raw data *M* into a genome-wide EpiCop matrix *P* ∈ *R^x^* ^×^ *^f^*, where *x* represents the total number of genomic regions. This matrix represents continuous EpiCop values indicating the strength of connection between genomic regions and EpiCops. To determine the association probabilistically, we applied a sigmoid function with a threshold of 0.5. Genomic regions with a probability above 0.5 were defined as being positively associated with a given EpiCop. This generates a genome-wide association matrix *A* that represents association of *i*-th genomic locus with *j*-th EpiCop.

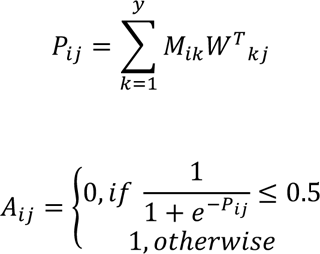

To analyse the genomic overlap between EpiCops, we first counted the number of 1kb windows co-occupied by a given pair of *i*-th (*A_i_*) and *j*-th EpiCops (*A*_j_). Then the Jaccard similarity index between this pair (*J_i_*_,j_) is defined as follows.

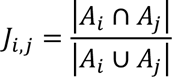

To perform pairwise comparisons of EpiCops, we computed cosine similarity for all EpiCop pairs using the cosine_similarity function from Python’s sklearn package. The weights from optimal RBM models across all 8 epigenetic marks were combined into a single matrix and z-scores normalized for each EpiCop were calculated. With this matrix, cosine similarity was calculated for each pair, with only values present in both EpiCops.

To extract EpiCop values for a given set of loci, we first concatenated the individual EpiCop matrices from the eight epigenetic marks into a single matrix. During concatenation, genomic loci lacking EpiCop values for a specific mark (e.g., due to absence of peaks at those loci) were imputed with zeros. The combined matrix was then normalised using z-scores. This concatenated matrix represents the continuous values of 720 EpiCops across genome-wide 1 kb windows. Genomic overlaps between input loci and EpiCop windows were assessed using bedtools intersect^60^. Loci were considered overlapping if at least one base pair overlapped. When a locus overlapped multiple EpiCop windows, it was assigned to the window with the greatest base-pair overlap.

To calculate tissue-level EpiCop values, we computed the average EpiCop z-score within each tissue group as defined by the EpiMap sample description^20^. For heatmap and UMAP visualisation, values were normalised across tissue groups for each EpiCop. Missing tissue-level values were imputed using the median value of the corresponding EpiCop.

### Functional characterisation of EpiCops using gene ontology (GO) and KEGG pathway terms

To characterise the functional scope captured by EpiCops, we first extracted protein-coding genes (PCGs) annotated with NM or XM RefSeq identifiers^26^. For each PCG, a 1 kb window was defined by extending 500 bp upstream and downstream of the transcription start site (TSS). Using a sigmoid activation function with a threshold of 0.5, we determined EpiCop associations across all PCGs. For each chromatin mark, EpiCops were ranked according to their contribution to genome-wide epigenomic variation using singular value decomposition (SVD) of the corresponding genomic EpiCop matrix. Contributions were calculated from the absolute EpiCop loadings across components explaining 95% of the total variance. The ten highest-ranking EpiCops for each chromatin mark were selected for downstream functional characterisation of protein-coding genes.

Within each epigenetic mark, a unique combination of these selected EpiCops was assigned to each PCG, generating groups of genes sharing distinct epigenomic patterns for that mark. This resulted in multiple pattern combinations per epigenetic mark. Only groups containing at least 10 genes were retained for downstream analysis. GO and KEGG enrichment analyses were performed for each PCG group using the g:Profiler Python package with default settings (one-sided Fisher’s exact test with Benjamini-Hochberg FDR correction)^61^. Enrichment analysis was conducted independently for each epigenetic mark to enable cross-mark comparison. GO and KEGG terms with FDR<0.05 were considered significant. For each epigenetic mark, we quantified the total number of non-redundant significant terms and identified terms shared across multiple pattern combinations, either within the same mark or across different marks. To assess term specificity, we calculated the proportion of EpiCop combinations associated with each enriched term. Lower proportions indicate greater specificity of a term to particular pattern combinations.

### Deep learning models for EpiCop-based genomic region classification

We extracted 35,625 enhancers across 51 in vivo tissue and cell types from EnhancerAtlas2.0 data resource. For each tissue and cell type, enhancers with hs score>1 are used. We also identified 2,386 pathogenic SNPs from 22 diseases, annotated as “Likely_pathogenic” or “Pathogenic/Likely_pathogenic” under the CLNSIG field, restricting the analysis to diseases with at least 100 pathogenic SNPs. For each enhancer or pathologic variant, overlapping 1kb windows are identified to extract EpiCop values in those regions. Z-scores were calculated for each EpiCop for the classification model training. For enhancers overlapping multiple 1kb windows, a single window that overlap the centre point of the corresponding enhancer was used.

We used deep neural network (DNN) models to predict associations between genomic regions based on their underlying epigenomic patterns. To identify optimal hyperparameters, we trained 10 DNN models with randomly sampled configurations. The hyperparameter search space included 4–8 hidden layers and 16, 32, 64, 128, 256, or 512 nodes per layer. Input data were represented as a pattern matrix *P* ∈ *R^x*720^*, where *x* denotes the number of genomic regions and the 720 columns correspond to individual EpiCops derived from eight epigenetic marks. Each region was labelled as positive or control for binary classification. The dataset was split into training and test sets at an 80:20 ratio, and missing values were imputed using the median. To address class imbalance, the synthetic minority over-sampling technique (SMOTE) was applied to the training set only^62^. Values were then normalized using z-scores. To improve generalization, we added an L2 ridge regularisation (*λ* = 0.001) and a dropout layer (with a probability of drop-out=0.5) applied to each hidden layer. Model performance was evaluated on the test set, and the configuration achieving the highest area under the ROC curve (AUC) was selected as optimal. Hidden layers used ReLU activation, the output layer used sigmoid activation, and models were trained with the Adam optimiser and binary cross-entropy loss (or categorical cross-entropy for multi-class classification tasks).

### Atrial fibrillation-associated EpiCop profiles and tissue representation

To characterise regulatory programs associated with atrial fibrillation (AF), EpiCop z-scores were extracted from 1-kb genomic windows overlapping AF-associated variants. For each EpiCop, z-scores were averaged across AF-associated variants, and EpiCops were ranked in descending order according to their mean score. The 30 EpiCops with the highest mean scores were retained for visualisation. Bars represent the mean EpiCop z-score across AF-associated variants, with error bars indicating the standard error of the mean across variants.

To examine the tissue representation of these AF-associated EpiCops, we used an EpiCop-by-tissue score matrix describing the relative representation of each EpiCop across tissue groups. Tissue scores for the 30 highest-ranked AF-associated EpiCops were extracted directly from this matrix. For each EpiCop, the heart score was compared with the mean score across all non-heart tissue groups. The 95% confidence interval for the non-heart reference was calculated as the non-heart mean ±1.96*s.e.m., where the s.e.m. was calculated from the standard deviation of scores across non-heart tissue groups divided by the square root of the number of non-heart tissue groups.

### Enrichment of AF-associated variants in selected EpiCops

Genome-wide EpiCop-positive regions were defined as 1-kb windows with sigmoid-transformed EpiCop values >0.5. The union of regions positive for any H3K4me1-derived EpiCop was used as the H3K4me1 genomic background. The enriched EpiCops with the highest mean z-scores across AF-associated variants were selected, and enrichment of AF-associated windows within regions represented by these EpiCops was assessed relative to the remaining H3K4me1-positive genomic background using a 2*2 contingency table. Odds ratios and one-sided p-values were calculated using Fisher’s exact test. 95% confidence intervals for odds ratios were estimated on the log scale using the standard error of the log odds ratio; a 0.5 continuity correction was applied when any contingency-table cell contained zero observations.

### Calculation of EpiCop tissue scores

Variants were mapped to their corresponding 1-kb genomic windows, and EpiCop z-scores for these windows were extracted from the genome-wide window-by-EpiCop matrix. Tissue scores were then calculated by multiplying the resulting variant-by-EpiCop matrix by an EpiCop-by-tissue matrix containing normalized average tissue weights for each EpiCop. This yielded a variant-by-tissue score matrix, in which each tissue score represents the weighted sum of EpiCop z-scores according to their relative representation in that tissue. For analyses of variant sets, scores were averaged across variants for each tissue group.

### Composite data analysis

In composite data analysis, we aimed to assess the capacity of EpiCop profiles to segregate genomic loci representing distinct biological contexts. To this end, genomic loci from multiple biological groups were combined into a common input dataset. Three data resources were analysed: (1) five tissue-specific enhancer groups (lung, cerebellum, liver, left ventricle and pancreas) from EnhancerAtlas 2.0^34^, 2. 14 disease groups from the Global Biobank Meta-analysis Initiative (GBMI) (asthma, chronic obstructive pulmonary disease, heart failure, stroke, gout, venous thromboembolism, primary open-angle glaucoma, abdominal aortic aneurysm, idiopathic pulmonary fibrosis, thyroid cancer, hypertrophic cardiomyopathy, uterine cancer, acute appendicitis and appendectomy)^33^, and 3. Five groups of tissue-specific GTEx eQTLs (heart left ventricle, thyroid, kidney cortex, brain cortex, muscle skeletal)^35^. Loci assigned to more than one biological group were excluded to ensure that each locus represented a single tissue or disease context. For each data resource, the remaining loci were clustered according to their EpiCop profiles. To quantify enrichment of each known biological context within the resulting clusters, we calculated the observed proportion of loci belonging to each enhancer tissue, disease or GTEx tissue group within each cluster. Statistical enrichment was assessed against a null distribution generated from 1,000 random permutations of locus cluster assignments. For each cluster-group combination, the observed proportion was compared with the corresponding permutation-derived distribution to calculate an enrichment z-score. Clustering concordance with the known disease or tissue labels was quantified using adjusted Rand index (ARI), which measures agreement between two partitions while correcting for chance and does not require the partitions to contain the same number of clusters. Because EpiCop clustering can subdivide loci sharing the same biological label into multiple regulatory clusters, ARI was interpreted as a measure of overall correspondence rather than exact recovery of the predefined labels. Statistical significance was assessed against an empirical null distribution generated from 1,000 random permutations of locus labels.

For tissue-level analysis of GBMI disease-associated variants, we quantified whether variant-associated tissue scores were shifted relative to their corresponding genome-wide distributions. For each disease, epigenetic mark, and tissue, we first centered the tissue score of each input variant relative to the genome-wide median score for the corresponding epigenetic mark and tissue.

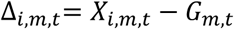

where *X_i_*_,*m*,*t*_ is a value for input variant *i*, epigenetic mark *m*, tissue *t*, and *G_m_*_,*t*_ is the median score across genome-wide windows for the corresponding epigenetic mark and tissue. The genome-wide median was used as the reference because genome-wide tissue-score distributions were highly skewed and the median provided a robust measure of central tendency. For each disease, the distribution of Δ*_i_*_,*m*,*t*_ values was then summarized across all associated variants separately for each epigenetic mark and tissue. We calculated the mean difference and its 95% confidence interval (i.e. mean±1.96*SE) where SE is the standard error of the mean.

For composite disease analysis, phenotypes were generated by combining two distinct phenotypes into a single cohort as described previously^63^. Composite phenotypes were comprised of asthma and gout, asthma and COPD, asthma and high blood pressure (or hypertension), or Type-1 diabetes and Type-2 diabetes. For each composite disease pair, the cohort in the UKBB was randomly split into a testing and training cohort across 10 permutations. For each permutation, we conducted a GWAS of the composite phenotype in the training cohort, using age, sex, Townsend deprivation factor and the first 10 principal components as covariates. Performance was evaluated for the top 100, 250, 500 and 1000 most significant, independent variants associated with the composite phenotype.

Lead variants were selected by linkage disequilibrium (LD) clumping using a P-value threshold of 5 ∗ 10^−8^. Variants were clustered using their EpiCop profiles using a consensus clustering procedure described previously^30^. Only genomic loci uniquely associated with a single group were included in the clustering analysis. Model parameters were varied iteratively, and the optimal configuration was selected based on R-squared value.

For each EpiCop-defined variant cluster, a partitioned polygenic score (pPGS) was calculated as the sum of effect-allele dosages weighted by the corresponding composite-phenotype GWAS effect estimates from the training cohort. An unpartitioned PGS comprising all variants included in the partitioned scores served as the comparator. All scores were standardised to a mean of zero and a standard deviation of one. Among cases within each composite cohort, cluster-specific pPGS were fitted jointly against individual disease subtype status. Predictive performance was quantified using Nagelkerke’s R-squared value, and incremental performance was calculated relative to a covariate-only model containing age, sex, Townsend deprivation index and the first 10 genetic principal components. Variability in performance was assessed across the 10 training-testing splits.

To assess differential disease associations, disease-specific pPGS coefficients from the test set and their standard errors were extracted from each EpiCop-defined cluster within each disease pair. For each matched cluster, the coefficient difference was calculated as

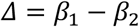

Where *β*_1_ and *β*_2_ represent effect size for disease 1 and 2 in each disease pair, respectively. Effect divergence was summarised as the perpendicular distance from the equality line (i.e. *β*_1_ = *β*_2_).

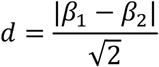

To quantify uncertainty of a measured difference, we used a conservative upper bound for its standard error because the covariance between the two estimates was unavailable. For coefficients *β*_1_ and *β*_2_, the variance of

their difference is SE_1_^2^ + SE_2_^2^ − 2*ρ*SE_1_SE_2_, where *ρ* denotes the correlation between the estimates. The maximum possible variance occurs at *ρ* = −1, yielding an upper-bound standard error of SE_1_ + SE_2_. A conservative z-statistic was therefore calculated as

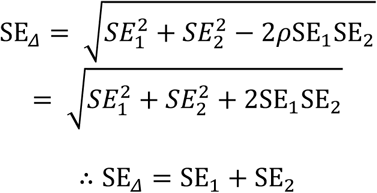

Then, two-sided approximate p-values were calculated from the statistic below.

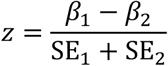

using the standard normal distribution. P-values were adjusted using the Benjamini–Hochberg procedure across all matched clusters within each disease pair. Clusters with adjusted p<0.05 were classified as having a detected differential effect, and their proportion was calculated relative to all matched clusters in that pair.

For the asthma-gout tissue analysis, clusters were selected using nominal p<0.05 in the disease-subtype model and assigned to asthma or gout according to negative or positive test coefficients, respectively, consistent with the phenotype coding. Remaining variants were labelled “Shared”. For each assigned disease group, variant-level EpiCop values were aggregated into average tissue scores and rescaled by min-max normalisation to the range 0-1, producing an epigenetic-mark-by-tissue matrix. The asthma matrix was subtracted from the gout matrix to summarise relative tissue patterns, with positive and negative differences indicating higher normalised scores in the gout and asthma groups, respectively.

### Clustering of type 2 diabetes-associated variant loci using EpiCops

Independent type 2 diabetes (T2D) genome-wide significant variants were selected from a published data^15^ with 643 SNPs included after filtering for overlap with available genomic annotations. For asthma, 419 SNPs were included after filtering^44^.

To perform clustering of genomic loci, we adopted a consensus clustering approach as previously described^30^. Note we modified parameters of this method, so that it fits better for our data. For instance, to reduce dimensionality of data, rather than using top 10 principal components, we used selected patterns (by median absolute deviation (MAD) value above 50^th^ percentile) to calculate k-nearest neighbours (k=15) in UMAP dimensions, using Euclidean distance. Subsequently, we performed Leiden community-based clustering with 100 different resolutions ranging from 0.2 to 20. By counting the number of times any pairs of genomic loci clustered to the same group, we created a consensus matrix. Linkage matrix was created with the corresponding distance matrix (calculated as 100 -consensus matrix), with Euclidean distance and Ward method. We iteratively performed hierarchical clustering with a varying number of clusters ranging from 2 to 300. At each iteration, we calculated a Silhouette score to quantify distinctiveness of variant clusters. By plotting a series of Silhouette scores, we found an optimal number of clusters at the minimum number of clusters where the average of preceding 5 Silhouette scores does not improve more than 0.001 - which effectively marked the start of plateau in many cases.

### Quantifying cluster specificity of EpiCops in T2D

To understand how specifically a given EpiCop is associated with T2D variant clusters, we used a Shannon entropy-based approach. For each i-th EpiCop, for its cluster-level profiles across the 72 T2D clusters, {*x*_1_, *x*_2_, …, *x*_72_ }, we computed 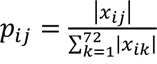, where *x_ij_* and *p_ij_* represent i-th EpiCop z-score and its proportional strength in j-th cluster, respectively. Subsequently, we computed normalised Shannon entropy, with 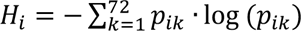 *p_ik_* · log (*p_ik_*). This was subsequently normalized by log(72), which represents the maximum H, i.e. 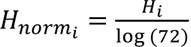. Finally, the specificity is calculated as 1-*H_norm_*, where 0 indicates broadly shared across clusters and 1 indicates maximal cluster specificity.

### Summary statistics-based cluster association with phenome

For T2D, we assembled a panel of clinically and biologically relevant traits spanning: (1) glucose homeostasis (2) anthropometric measures; (3) blood biomarkers; (4) lipid metabolism; (5) liver function (6) renal function; and (7) cardiovascular traits. For asthma, the phenotype panel included (1) blood biomarkers; (2) immune cell populations; (3) respiratory function measures; and (4) asthma subtypes. GWAS summary statistics for these phenotypes were obtained from multiple consortia including UK Biobank, MAGIC, GIANT, and GWAS catalogues^64–67^. All GWAS summary statistics were harmonised such that effect alleles were aligned to the disease risk-increasing alleles. For T2D, effect alleles were oriented to match the T2D risk allele from the reference GWAS ^16^. For asthma, effects were similarly aligned to the asthma risk allele from Stikker et al.^44^.

For each EpiCop cluster and phenotype, we aggregated variant-level GWAS effect estimates using inverse-variance weighted fixed-effects meta-analysis. This approach weights each variant by the precision of its effect estimate. The resulting summary estimate was defined as the aggregated effect size (AES) which represents the overall direction and magnitude of association between variants in a given cluster and the phenotype. We also included an all variant set containing all disease-associated variants as a control. Only non-redundant phenotypes were retained for enrichment analysis. Cluster–phenotype tests were excluded when no variants from the cluster overlapped the corresponding GWAS summary statistics file.

To assess enrichment of phenotypes among variant clusters compared to random association, we created a null distribution of enrichment derived from 1,000 random shuffling permutations. We calculated proportions of variants associated with a given phenotype in each cluster, so that the sum of the proportions of that phenotype across clusters is 1. Subsequently, z-scores of the proportion were used to identify significantly enriched cluster-phenotype pairs (FDR<0.05, two-sided z-test). In addition, we adopted adjusted rand index (ARI) to assess the overall clustering agreement by comparing against ground-truth of their associations if known (e.g. tissue-type specific enhancers). To calculate statistical significance, ARI values derived from 1,000 random shuffling permutations were used as the null distribution with an assumption of normal distribution (t-test, two-sided).

### Functional Correlation between variant clusters

To quantify the distinctiveness of each cluster’s phenotype association profile relative to the global average, we calculated deviation scores. For each phenotype k and cluster j, the deviation was computed as: *D*_j*k*_ = *AES*_j*k*_ − *AES*_0*k*_, where *AES*_j*k*_ is the aggregated effect size for cluster *jj*on phenotype *k*, and *AES*_0*k*_ is the aggregated effect size for the all variant whole genome control on phenotype *k*. The deviation profile for cluster *jj* can therefore be written as D_j_ = (*D*_j1_, *D*_j2_, …, *D*_jK_).

For each Smith-defined T2D cluster *x* and each EpiCop-defined T2D cluster *y*, the functional similarity was then quantified using Spearman correlation between their deviation profiles 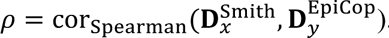.

### UK Biobank and All of US individual-level analyses

#### Population cohorts

UK Biobank^64^ is a prospective population-based cohort of approximately 500,000 UK participants recruited at ages 40–69, with linked genetic, lifestyle, biomarker, hospital, primary care, medication, and mortality data.

All of Us^51^ is a US-based population-based cohort integrating participant surveys, physical measurements, electronic health records, wearable-device data, and genomic data; the v8 release includes genomic data for more than 414,000 with short-read whole-genome sequencing.

#### Genotype pre-processing

UK Biobank imputed genotype data were processed using PLINK 2.0^68^. Genotypes were restricted to biallelic variants, and variatns with minor allele frequency (MAF)<0.01, Hardy-Weinberg equilibrium P<1×10⁻⁶, or imputation quality score (INFO)<0.7 were excluded. After quality control, genotype data from chromosomes 1-22 were merged across all autosomes. The merged genotype dataset was restricted to European unrelated individuals.

We used All of Us ACAF-threshold genotype from v8 release. Genotypes were pre-processed using PLINK 2.0, restricted to biallelic variants, and filtered to retain unrelated European-ancestry individuals.

#### Cohort and covariates definitions

Type 2 diabetes cases were defined using the eMERGE Type 2 Diabetes Mellitus phenotype algorithm^69^, which classifies cases based on combinations of T2D ICD9/10 diagnosis codes diabetes medication exposures, and supporting laboratory evidence, while excluding individuals with type 1 diabetes diagnosis codes. Controls were defined as participants with no diabetes-related diagnostic, self-report, medication, insulin, or laboratory evidence.

Asthma cases were defined as individuals with ICD9/10 diagnosis codes as well as those with self-reported asthma. Controls were defined as individuals without any asthma-related diagnosis codes. Controls were defined as participants with no asthma-related diagnosis codes and no self-reported asthma. Individuals with allergic or asthma-related conditions were excluded from the controls.

Covariates used include age, sex, Townsend deprivation index, the first 10 genetic principal components, disease tatus where relevant, and trait-specific medication covariates. Medication covariates included lipid medication for lipid traits, glucose medication for glycaemic traits, and hypertension medication for blood pressure traits.

#### Polygenic risk score calculation

T2D and asthma genome wide significant variants from Smith et al. and Stikker et al. were matched to variants in the UK Biobank imputation array. Variants not directly available were matched with LD proxies (r² > 0.7) using 1000 Genomes Project phase 3 European reference genotype^70^ Variant weights were additionally mapped to independent, non-UK Biobank GWAS cohorts to enable out-of-sample prediction as previously described^71^.

Baseline whole genome polygenic risk scores were calculated using all disease-associated variants weighted by their marginal GWAS effect sizes. For individual *i*, the whole-genome PGS was defined as:

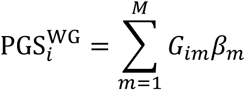

where *G_im_* ∈ {0,1,2} is the effect-allele dosage for variant *m*, *β_m_*is the GWAS effect size, and *M*is the number of disease-associated variants

Cluster partitioned PGS were calculated separately for each epigenomic cluster using only variants assigned to that cluster, weighted by their GWAS effect sizes using plink. For each epigenomic cluster *jj*, the cluster-partitioned PGS was defined as:

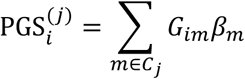

where *C*_j_ is the set of variants assigned to EpiCop cluster *jj*. All PGS were standardised before downstream modelling.

#### PGS model evaluation and association testing

Logistic and linear models were used respectively for binary and continuous phenotypes. Covariates include age at assessment, sex, the first 10 genetic principal components, Townsend deprivation index, disease-relevant medication use, and baseline disease status.

UKB analyses used train/test splits generated from the case-control cohort. For each split, PGS values were merged with covariates and phenotypes, and analyses were run separately for T2D-only participants and the full cohort with or without T2D status included as a covariate.

For each phenotype, three models were evaluated:

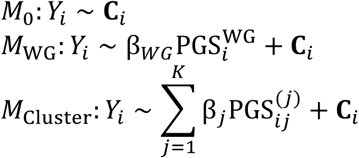

where *Y_i_*is the phenotype for individual *i*, C*_i_*is the covariate vector, 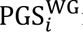 is the whole-genome or all-variant disease PGS, and 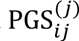 is the PGS for EpiCop cluster *jj* with coefficient β_j_. Binary outcomes were modelled using logistic regression and evaluated using Nagelkerke’s pseudo-*R*^2^. Continuous outcomes were modelled using linear regression and evaluated using *R*^2^. Models were fit in the training set and evaluated in the test set by regressing the observed phenotype on the model prediction.

The incremental variance explained by whole-genome and cluster-partitioned PGS was calculated as:

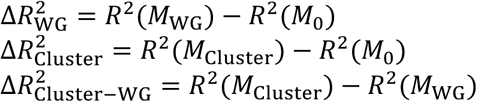

Permutation analyses were conducted over repeated random train/test splits to estimate the stability of model performance and the empirical distribution of *R*^2^gains.

In addition, we performed per-cluster association analyses to identify which individual EpiCop clusters were associated with specific phenotypes using the full cohort. For each phenotype, each cluster-specific PGS was tested in a separate regression model adjusted for the same covariates:

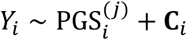

where 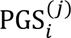 is the standardised PGS for cluster *jj*in individual *i*. For binary phenotypes, logistic regression was used with odds reported. For continuous phenotypes, linear regression was used with regression coefficients reported.

#### Cardiac incidence analysis

To test whether EpiCop-partitioned T2D genetic burden stratified cardiovascular outcomes, time-to-event analyses were performed using UKB follow-up data. Cluster-specific PGS were calculated for individual EpiCop clusters.

For myocardial infarction incidence analyses, prevalent myocardial infarction was removed by excluding individuals with MI dates before recruitment. Follow-up time was calculated from recruitment to the earliest of myocardial infarction or administrative censoring.

Individuals were stratified by cluster PGS quantiles. For single-cluster analyses, individuals in the top 30% and bottom 30% of the cluster-specific PGS distribution were compared. For cross-cluster analyses, discordant PGS groups were defined using clusters 10 and 36:

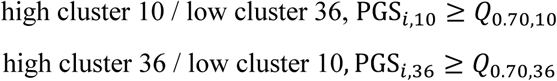

Cox proportional hazards models were fit as: 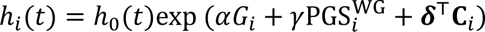 where ℎ*_i_*(*t*)is the hazard for individual *i*, ℎ_0_(*t*)is the baseline hazard, *G_i_* is the PGS group, 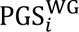 is the whole-genome T2D PGS, and C*_i_* includes age, sex, PCs 1–10, Townsend deprivation index, medication covariates, and T2D status.

### Consistency of phenotype association between variant clustering methods

To assess consistency of phenotypes captured by different clustering methods and parameters, we repeated clustering of T2D variants from Smith et al.^15^ using k-means, hierarchical and Leiden clustering method. Clustering parameters were tuned to generate the same number of variant clusters as obtained with a modified consensus clustering method^30^ (i.e. 72 clusters). Pairwise similarity between clustering results was quantified using the adjusted rand index (ARI).

### Cross-architecture sensitivity analysis with matched latent dimensionality

To assess whether an alternative latent-variable model recovered similar epigenomic structure at the same latent dimensionality, we trained a linearly decoded variational autoencoder (LD-VAE)^72^. A LD-VAE was trained per chromatin mark for comparison to the RBMs. The latent dimensionality for each chromatin mark was fixed to the optimised hidden-node count of the corresponding RBM, yielding 720 latent features across assays. The encoder and decoder used single-layer architectures with ReLU activation. Chromosomes 8 and 10 were reserved for validation, chromosomes 9 and 11 for testing, and the remaining chromosomes for training. The training objective combined binary cross-entropy reconstruction loss with Kullback–Leibler divergence regularisation. The learning rate followed a cosine-annealing schedule after linear warmup, and the KL-divergence weight was increased using a linear warmup schedule. The model checkpoint with the lowest validation loss was selected for testing and downstream analyses.

We compared the downstream clustering behaviour of the two representations using T2D-associated variants from Smith et al.^15^ All 720 features from each model were used for clustering, with resolution parameters adjusted to obtain 72 clusters for each representation. We additionally evaluated mean silhouette scores for candidate cluster counts ranging from 2 to 300 and compared the range of cluster-level aggregated effect-size enrichment between representations. This analysis assessed sensitivity to model architecture at matched latent dimensionality, rather than the relative performance of independently optimised architectures.

### Benchmarking analysis for variant clustering

We performed a head-to-head benchmarking analysis to compare EpiCop-derived clustering of T2D variants with existing approaches: 1. Clustering using disease-relevant phenotypes, as reported in Smith et al.^15^, 2. Direct epigenomic annotations, using the high confidence variant set only as previously described^44^, 3. Integrative epigenomic and pleotropic modelling^42^, using J-PEP W factorised matrix for T2D variant from Suzuki et al.^16^, 4. Tissue-specific open chromatin, using EpiMap DNase data^20^, 5. Genome-wide decomposition of 2,138 phenotypes ^43^. For clustering variants using J-PEP W matrix^42^, we used data for ‘All_Metal_LDSC-CORR_Neff_W’ and assigned T2D variants to a cluster with the highest score. For tissue-specific open chromatin data^20^, T2D variants were mapped to EpiMap tissue-enriched DNase-seq 1 kb windows by direct overlap or through LD proxies. We identified T2D variants from Smith et al. that overlapped with open chromatin loci in multiple tissues, including digestive, immune B and T cell, liver, lymphoid, muscle, musculoskeletal, neural, pancreatic, pluripotent, reproductive, respiratory, urinary and vascular systems, as derived from EpiMap DNase I hypersensitivity assays. Subsequently, we created a variant-by-tissue binary overlap matrix. We clustered variants based on their shared tissue-specificity profiles. We only kept variants clusters with at least 5 variants, yielding 13 clusters composed of 348 variants. To keep the variant set identical between methods, we used the same set of variants for EpiCop clustering. We used only EpiCops with MAD>0.5 for variant clustering. In this application, we clustered variants using Leiden clustering method, with the same cluster number (k=13) as tissue-specific open region clustering. For performance comparison, we used 22 non-redundant cardiometabolic traits. Variant clusters without any mapping variants from phenotype enrichment analysis were excluded from the benchmarking analysis. For genotype-phenotype genomic decomposition method (DeGAs)^43^, we assigned variants to a single feature (or PC) with the highest score, using ‘dev_allNonMHC_z_center_p0001_100PCs_20180129.npz’ file from the study. To this end, we first performed overlap analysis between their reported contribution variants and genome-wide EpiCop windows that overlap T2D variants from Suzuki et al. (as reported in Kerner et al.^42^), yielding 392 variants used for clustering analysis. We ran Leiden clustering with a resolution of 1.0 based on DeGAs 100 PCs, which gave 7 clusters. To get the same number of clusters with EpiCops using Leiden clustering, we used a resolution of 0.7, with a random seed of 42.

For tissue-specific open chromatin method, we first identified tissue specific open variants obtained from Suzuki et al. present in relevant tissue groups including EpiMap tissue annotations, digestive, immune B and T cell, liver, lymphoid, muscle, musculoskeletal, neural, pancreatic, pluripotent, reproductive, respiratory, urinary and vascular systems. Subsequently, we created a variant-by-tissue binary overlap matrix. We grouped variants sharing identical overlap binary profiles. We removed clusters with less than 5 variants, yielding a total of 13 clusters composed of 348 variants. We independently ran clustering of these variant using Leiden clustering method based on EpiCops, with the same cluster k number (k=13).

