## Supplementary Figures for "Epigenomic regulatory programs reveal the latent heterogeneity of complex disease"

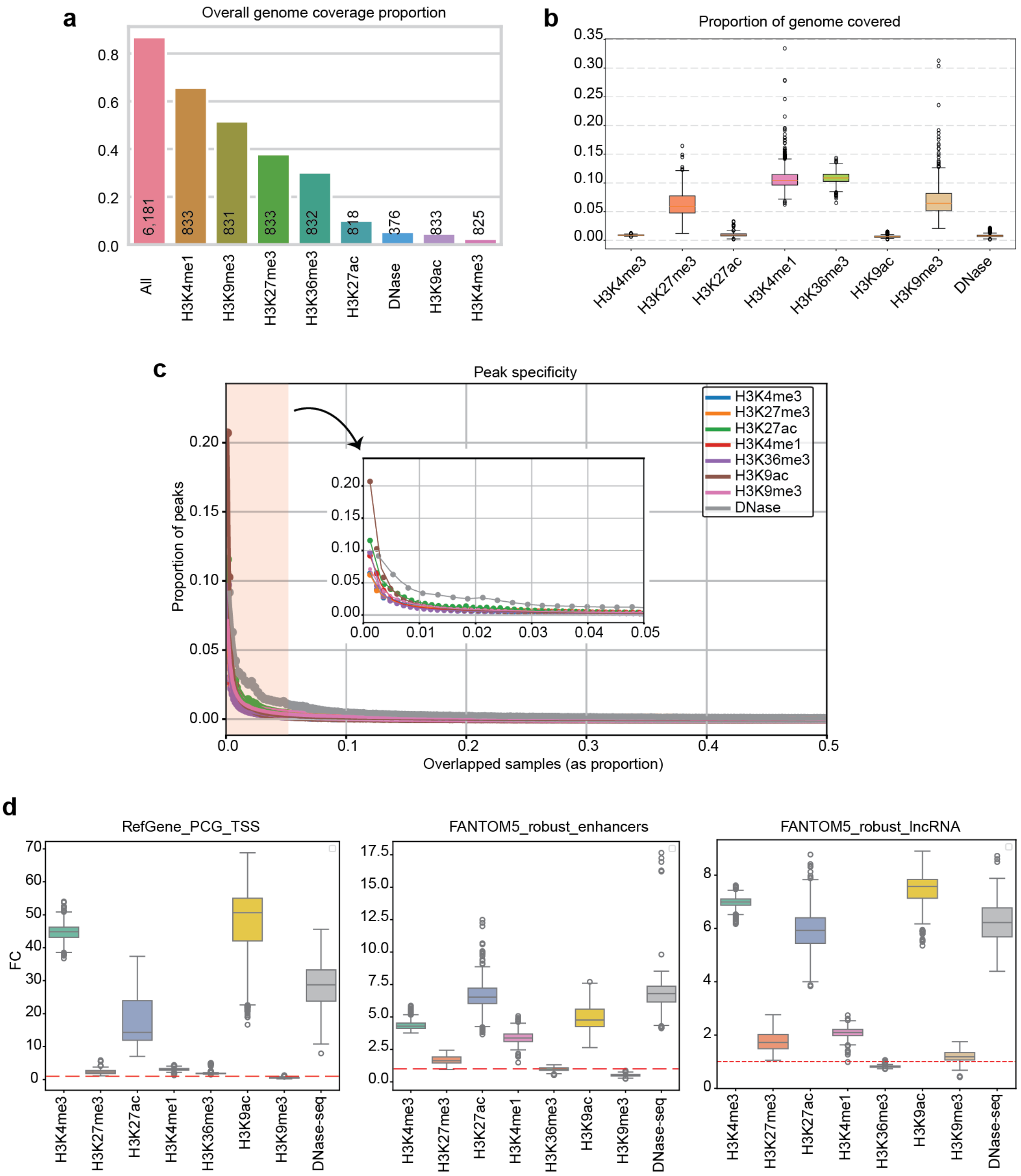


**Supplementary Figure 1.** EpiMap consortium epigenomic data for EpiCop extraction.

**a**. Proportion of the genome covered by the combined set of peaks. Numbers within the bars indicate the number of epigenomic profiles, with 'All' representing the combined total across chromatin modalities.

**b.** Proportion of the genome covered by each sample for each epigenetic mark.

**c**. Peak specificity, shown as the proportion of samples in which each peak was observed. For each epigenetic mark, all peaks were assessed for the number of samples where peaks were co-occupied. These values were then expressed as a proportion of the total number of peaks for that epigenetic mark.

**d**. Association of peaks with genomic regulatory elements. Fold enrichment was calculated relative to random genomic associations. FC of 1 indicates (red-dashed line) random association.

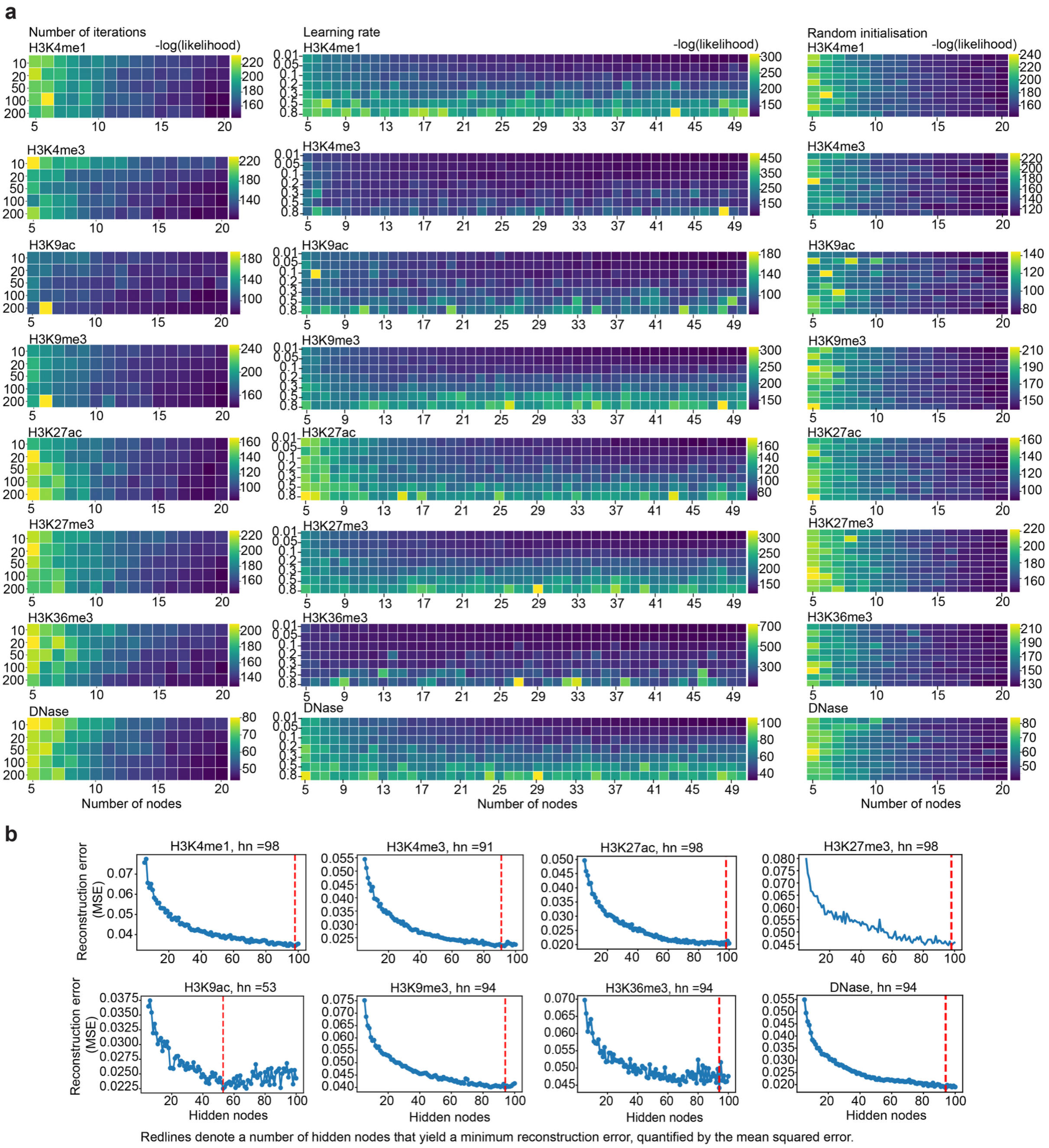


**Supplementary Figure 2.** RBM model training.

**a**. Hyperparameter selection. Values represent the mean log-likelihood across five-fold cross-validation.

**b**. The number of hidden nodes yielding the lowest generalisation error measured by mean squared error, highlighted with the red dashed line.

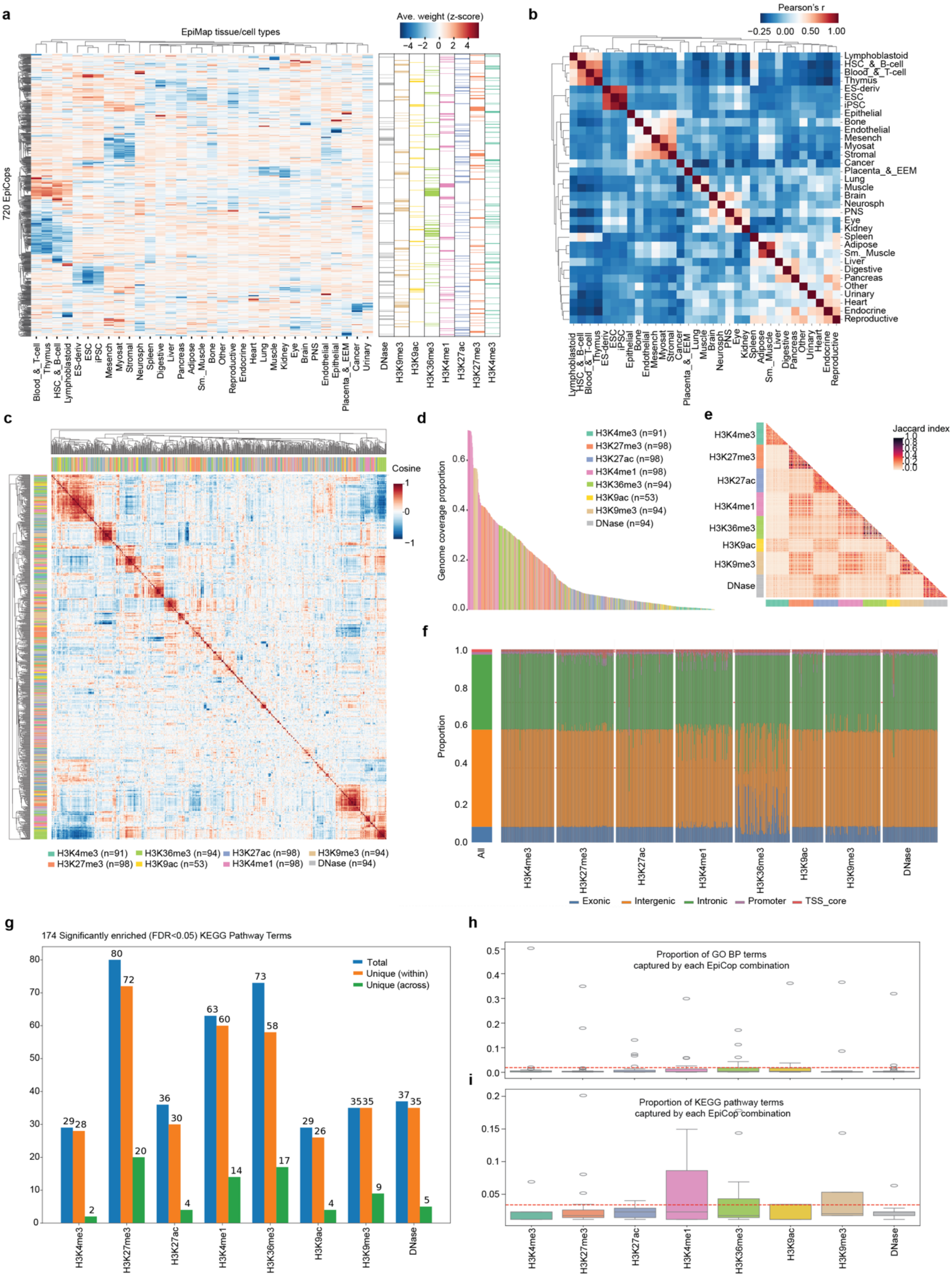


**Supplementary Figure 3.** Characterisation of EpiCops.

**a**. Tissue-level EpiCop weights.

**b**. Pearson correlation coefficients between EpiMap tissue groups based on EpiCop profiles.

**c**. Cosine similarity of EpiCop representations.

**d**. Genomic coverage of EpiCops

**e**. Genomic co-occurrence of EpiCops.

**f**. Genomic distribution of EpiCops.

**g**. Enrichment of KEGG pathway terms in genomic loci enriched with EpiCop combinations across chromatin marks (sigmoid probability>0.5).

**h-i**. Proportion of GO BP (h) and KEGG (i) terms captured by individual EpiCop combinations.

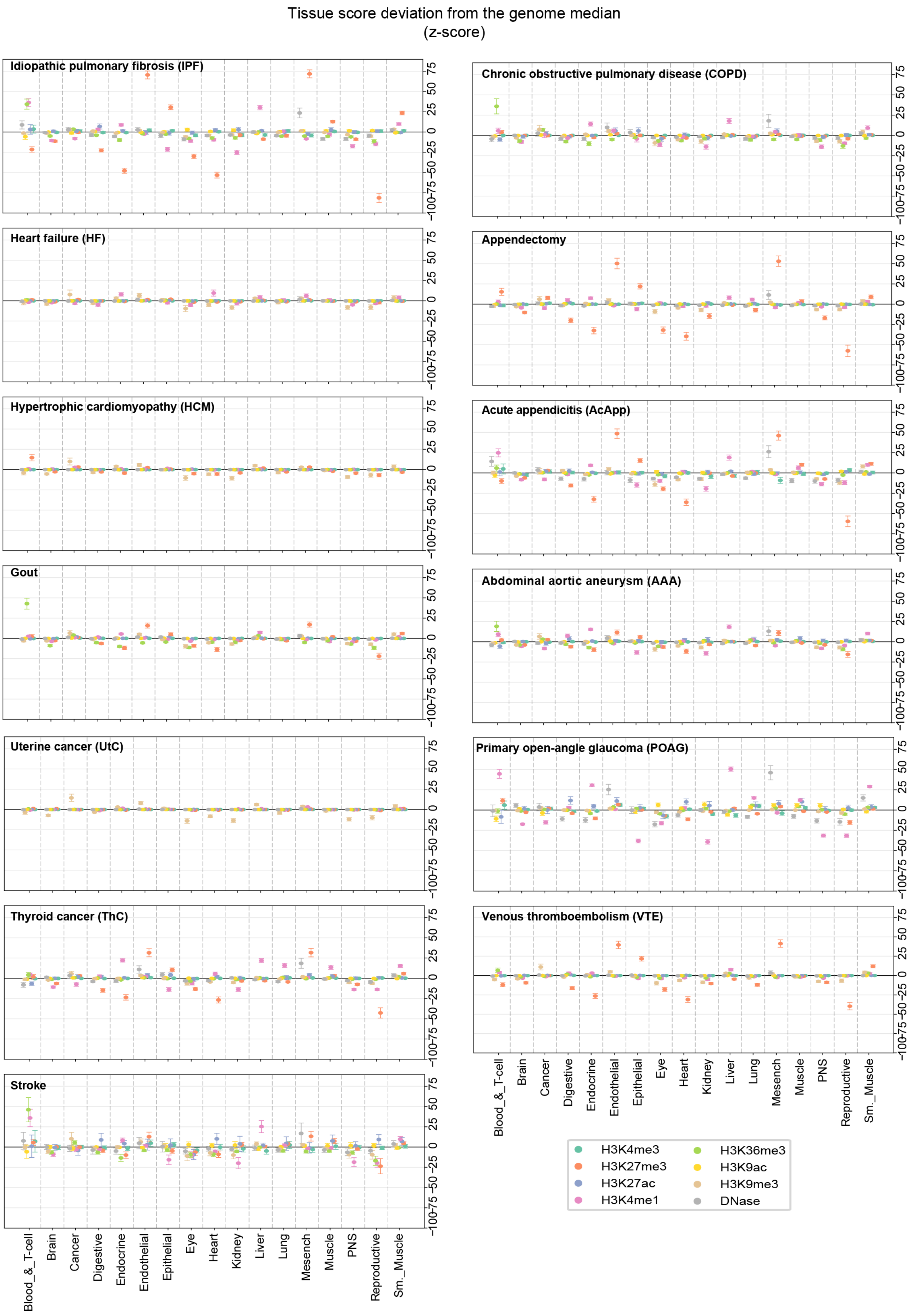


**Supplementary Figure 4.** Tissue-level epigenetic enrichments at disease variants across different chromatin marks. Values represent deviations of tissue-level EpiCop z-scores from the corresponding genome-wide medians.

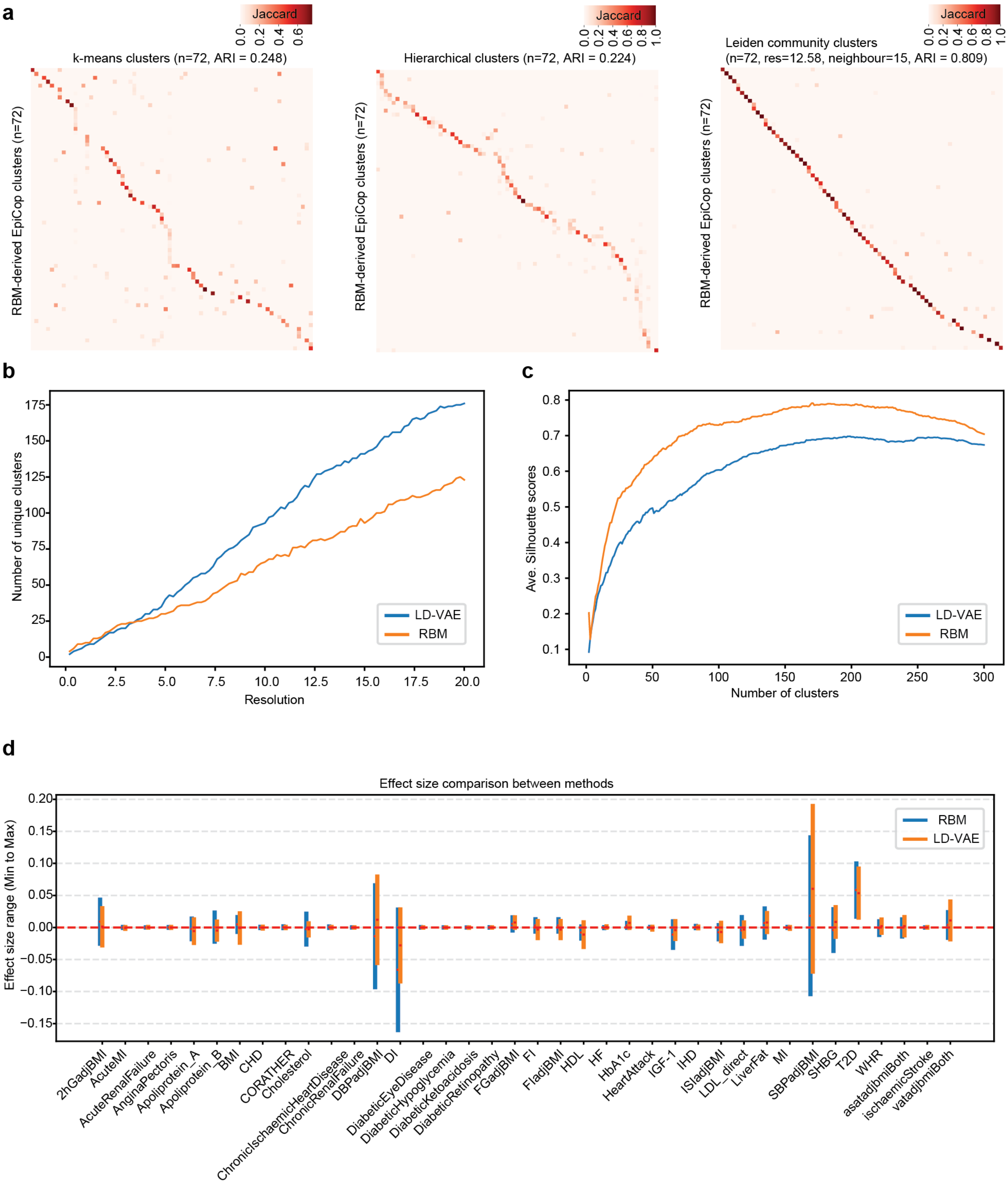


**Supplementary Figure 5.** Clustering consistency between model architectures using different clustering and pattern extraction algorithms.

**a**. Overlap of T2D variants across clusters defined by the original consensus clustering approach and three common clustering algorithms (i.e. k-means, hierarchical or Leiden clustering), with the number of clusters matched to the original. Colour indicates Jaccard similarity between clusters.

**b**. Number of T2D clusters identified by EpiCops derived from two different approaches (RBM vs LD-VAE) across Leiden clustering resolutions.

**c**. Cluster quality assessed by average silhouette score across varying numbers of clusters.

**d**. Head-to-head comparison between RBM- and LD-VAE-derived EpiCops for detecting enrichment of cardiometabolic traits, with the optimal number of clusters used for each method.

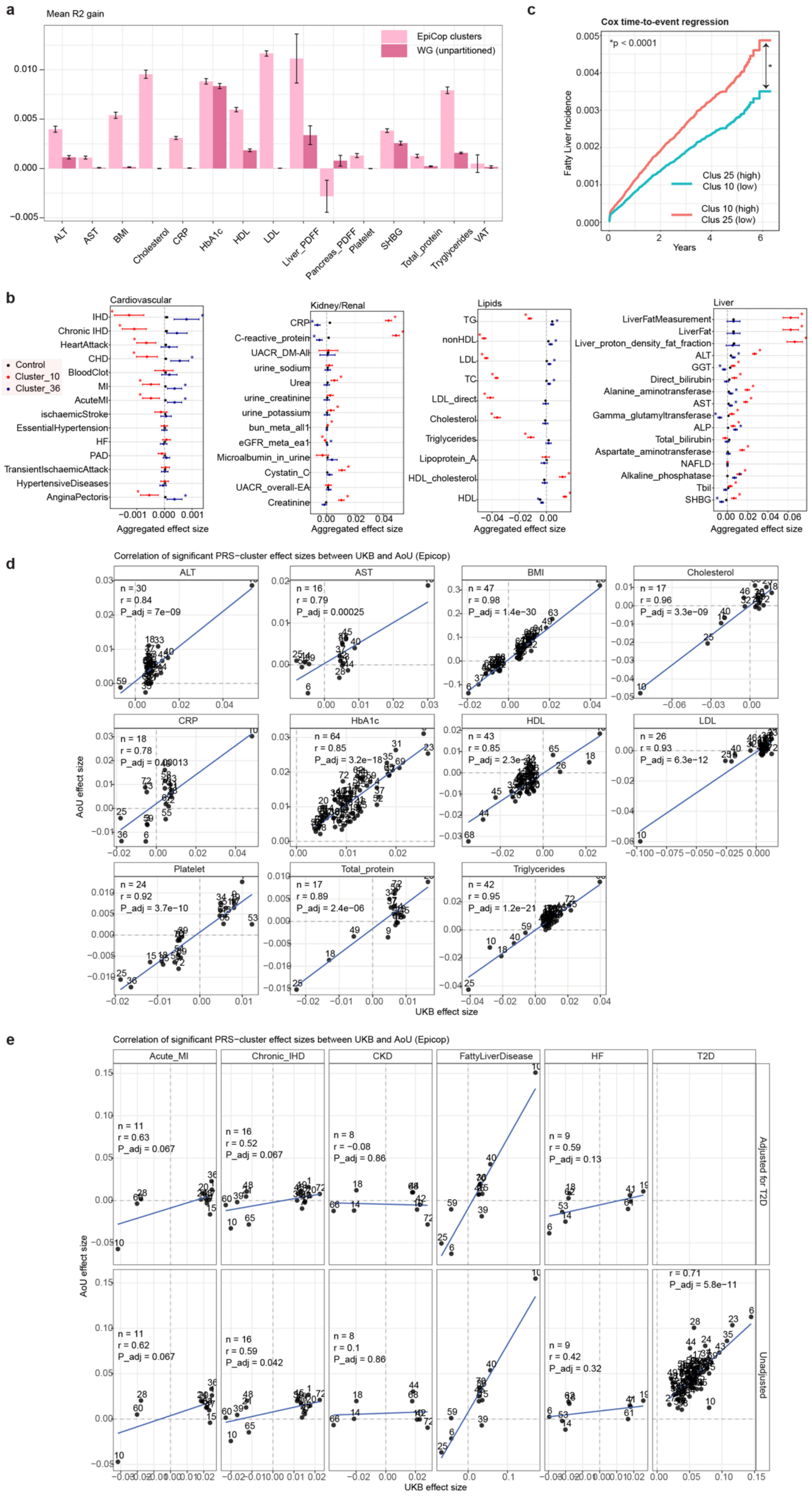
a
Mean R2 gain
EpiCop clusters
WG (unpartitioned)
ALT
AST
BMI
Cholesterol
CRP
HbA1c
HDL
LDL
Liver_PDFF
Pancreas_PDFF
Platelet
SHBG
Total_protein
Tryglycerides
VAT
c
Cox time-to-event regression
*p < 0.0001
Fatty Liver Incidence
Clus 25 (high)
Clus 10 (low)
Clus 10 (high)
Clus 25 (low)
Years
b
Cardiovascular
Control
Cluster_10
Cluster_36
IHD
Chronic IHD
HeartAttack
CHD
BloodClot
MI
AcuteMI
ischaemicStroke
EssentialHypertension
HF
PAD
TransientIschaemicAttack
HypertensiveDiseases
AnginaPectoris
Aggregated effect size
Kidney/Renal
CRP
C-reactive_protein
UACR_DM-All
urine_sodium
Urea
urine_creatinine
urine_potassium
bun_meta_all1
eGFR_meta_ea1
Microalbumin_in_urine
Cystatin_C
UACR_overall-EA
Creatinine
Aggregated effect size
Lipids
TG
nonHDL
LDL
TC
LDL_direct
Cholesterol
Triglycerides
Lipoprotein_A
HDL_cholesterol
HDL
Aggregated effect size
Liver
LiverFatMeasurement
LiverFat
Liver_proton_density_fat_fraction
ALT
GGT
Direct_bilirubin
Alanine_aminotransferase
AST
Gamma_glutamyltransferase
ALP
Total_bilirubin
Aspartate_aminotransferase
NAFLD
Alkaline_phosphatase
Tbil
SHBG
Aggregated effect size
d
Correlation of significant PRS−cluster effect sizes between UKB and AoU (Epicop)
ALT n = 30 r = 0.84 P_adj = 7e−09
AST n = 16 r = 0.79 P_adj = 0.00025
BMI n = 47 r = 0.98 P_adj = 1.4e−30
Cholesterol n = 17 r = 0.96 P_adj = 3.3e−09
CRP n = 18 r = 0.78 P_adj = 0.00013
HbA1c n = 64 r = 0.85 P_adj = 3.2e−18
HDL n = 43 r = 0.85 P_adj = 2.3e−
LDL n = 26 r = 0.93 P_adj = 6.3e−12
Platelet n = 24 r = 0.92 P_adj = 3.7e−10
Total_protein n = 17 r = 0.89 P_adj = 2.4e−06
Triglycerides n = 42 r = 0.95 P_adj = 1.2e−21
AoU effect size
UKB effect size
e
Correlation of significant PRS−cluster effect sizes between UKB and AoU (Epicop)
Acute_MI
Chronic_IHD
CKD
FattyLiverDisease
HF
T2D
Adjusted for T2D
n = 11 r = 0.63 P_adj = 0.067
n = 16 r = 0.52 P_adj = 0.067
n = 8 r = −0.08 P_adj = 0.86
n = 9 r = 0.59 P_adj = 0.13
Unadjusted
n = 11 r = 0.62 P_adj = 0.067
n = 16 r = 0.59 P_adj = 0.042
n = 8 r = 0.1 P_adj = 0.86
n = 9 r = 0.42 P_adj = 0.32
r = 0.71 P_adj = 5.8e−11
AoU effect size
UKB effect size

**Supplementary Figure 6. EpiCop-derived T2D-associated variant clusters.**

**a.** Comparison of mean R-squared gain for T2D-related biochemical traits between polygenic models partitioned by EpiCop clusters and the genome-wide (WG) unpartitioned model. Bars indicate R-squared gain, with error bars representing the standard error or the mean.

**b.** Meta-analysis of cardiometabolic trait enrichment in T2D variant clusters 10 and 36. Asterisks indicate FDR<0.05.

**c**. Cox time-to-event analysis of incident fatty liver disease in individuals stratified by EpiCop clusters 10 and 25.

**d-e**. Pearsons's correlation coefficients of effect sizes for significant cluster-specific PRS associations for continuous (**d**) or binary (**e**) cardiometabolic traits between UK Biobank (UKB) and All of Us (AoU) cohorts. The blue line represents a least-squares regression fit for each trait. Each point corresponds to a cluster defined in UKB, with the corresponding cluster identification number shown.

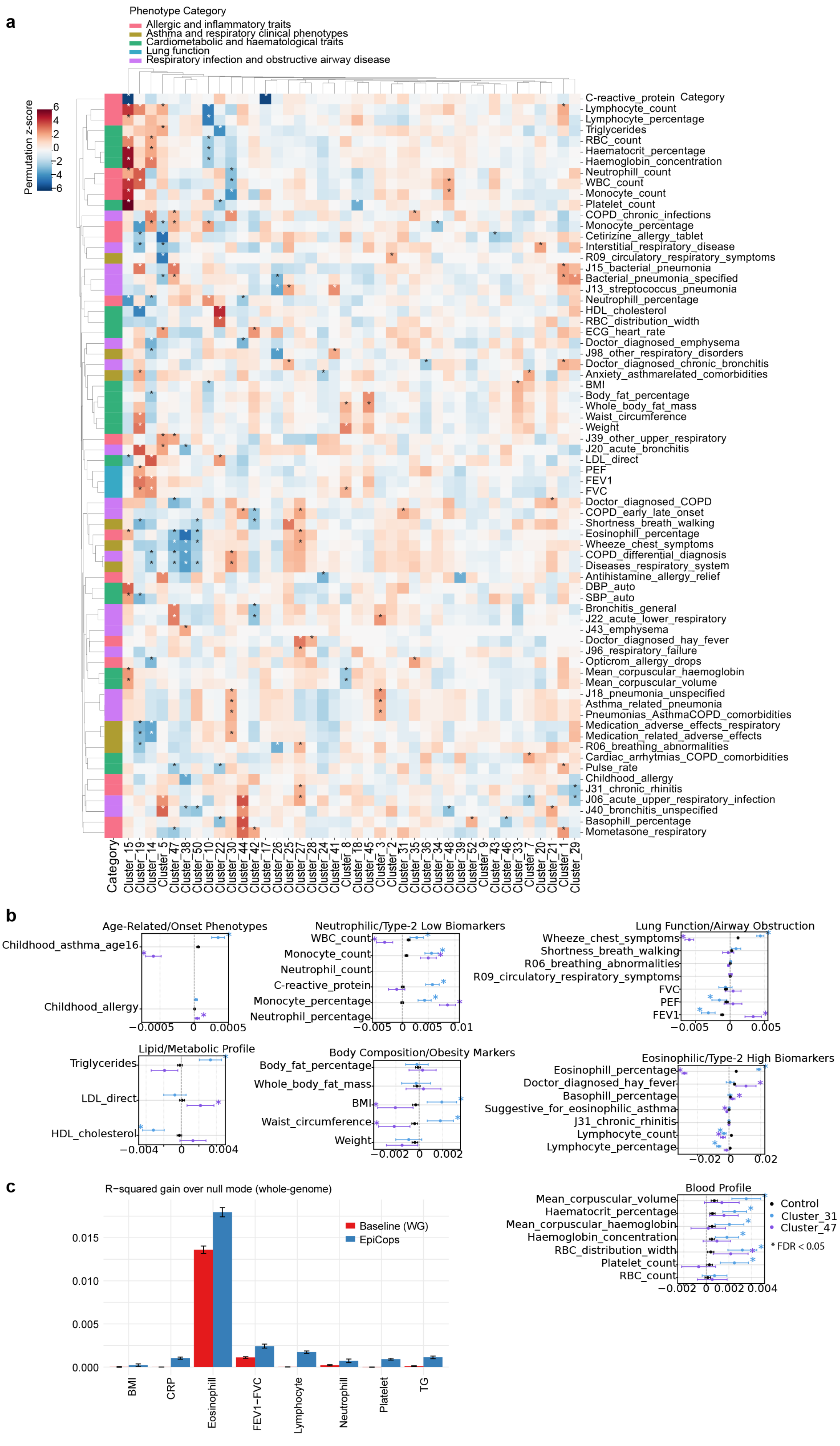

a
Phenotype Category
Allergic and inflammatory traits
Asthma and respiratory clinical phenotypes
Cardiometabolic and haematological traits
Lung function
Respiratory infection and obstructive airway disease
Permutation z-score
6
4
2
0
-2
-4
-6
Category
C-reactive_protein
Lymphocyte_count
Lymphocyte_percentage
Triglycerides
RBC_count
Haematocrit_percentage
Haemoglobin_concentration
Neutrophill_count
WBC_count
Monocyte_count
Platelet_count
COPD_chronic_infections
Monocyte_percentage
Cetirizine_allergy_tablet
Interstitial_respiratory_disease
R09_circulatory_respiratory_symptoms
J15_bacterial_pneumonia
Bacterial_pneumonia_specified
J13_streptococcus_pneumonia
Neutrophill_percentage
HDL_cholesterol
RBC_distribution_width
ECG_heart_rate
Doctor_diagnosed_emphysema
J98_other_respiratory_disorders
Doctor_diagnosed_chronic_bronchitis
Anxiety_asthmarelated_comorbidities
BMI
Body_fat_percentage
Whole_body_fat_mass
Waist_circumference
Weight
J39_other_upper_respiratory
J20_acute_bronchitis
LDL_direct
PEF
FEV1
FVC
Doctor_diagnosed_COPD
COPD_early_late_onset
Shortness_breath_walking
Eosinophill_percentage
Wheeze_chest_symptoms
COPD_differential_diagnosis
Diseases_respiratory_system
Antihistamine_allergy_relief
DBP_auto
SBP_auto
Bronchitis_general
J22_acute_lower_respiratory
J43_emphysema
Doctor_diagnosed_hay_fever
J96_respiratory_failure
Opticrom_allergy_drops
Mean_corpuscular_haemoglobin
Mean_corpuscular_volume
J18_pneumonia_unspecified
Asthma_related_pneumonia
Pneumonias_AsthmaCOPD_comorbidities
Medication_adverse_effects_respiratory
Medication_related_adverse_effects
R06_breathing_abnormalities
Cardiac_arrhytmias_COPD_comorbidities
Pulse_rate
Childhood_allergy
J31_chronic_rhinitis
J06_acute_upper_respiratory_infection
J40_bronchitis_unspecified
Basophill_percentage
Mometasone_respiratory
Category
Cluster_15
Cluster_19
Cluster_14
Cluster_5
Cluster_47
Cluster_38
Cluster_50
Cluster_10
Cluster_22
Cluster_30
Cluster_44
Cluster_42
Cluster_17
Cluster_26
Cluster_25
Cluster_27
Cluster_28
Cluster_24
Cluster_41
Cluster_8
Cluster_18
Cluster_45
Cluster_3
Cluster_2
Cluster_31
Cluster_35
Cluster_36
Cluster_34
Cluster_48
Cluster_39
Cluster_52
Cluster_9
Cluster_43
Cluster_46
Cluster_33
Cluster_7
Cluster_20
Cluster_21
Cluster_1
Cluster_29
b
Age-Related/Onset Phenotypes
Childhood_asthma_age16
Childhood_allergy
−0.0005
0
0.0005
Neutrophilic/Type-2 Low Biomarkers
WBC_count
Monocyte_count
Neutrophil_count
C-reactive_protein
Monocyte_percentage
Neutrophil_percentage
−0.005
0
0.005
0.01
Lung Function/Airway Obstruction
Wheeze_chest_symptoms
Shortness_breath_walking
R06_breathing_abnormalities
R09_circulatory_respiratory_symptoms
FVC
PEF
FEV1
−0.005
0
0.005
Lipid/Metabolic Profile
Triglycerides
LDL_direct
HDL_cholesterol
−0.004
0
0.004
Body Composition/Obesity Markers
Body_fat_percentage
Whole_body_fat_mass
BMI
Waist_circumference
Weight
−0.002
0
0.002
Eosinophilic/Type-2 High Biomarkers
Eosinophill_percentage
Doctor_diagnosed_hay_fever
Basophill_percentage
Suggestive_for_eosinophilic_asthma
J31_chronic_rhinitis
Lymphocyte_count
Lymphocyte_percentage
−0.02
0
0.02
Blood Profile
Mean_corpuscular_volume
Haematocrit_percentage
Mean_corpuscular_haemoglobin
Haemoglobin_concentration
RBC_distribution_width
Platelet_count
RBC_count
−0.002
0
0.002
0.004
Control
Cluster_31
Cluster_47
* FDR < 0.05
c
R−squared gain over null mode (whole-genome)
Baseline (WG)
EpiCops
0.015
0.010
0.005
0.000
BMI
CRP
Eosinophill
FEV1−FVC
Lymphocyte
Neutrophill
Platelet
TG

**Supplementary Figure 7.** Clustering of asthma-associated variants using the EpiCop-derived regulatory programs.

**a**. Enrichment of asthma-relevant phenotypes across EpiCop-based clusters. Asterisks indicate absolute z-score above 2 compared with a null distribution obtained from 1,000 random permutations.

**b**. Meta-analysis comparing the effect sizes of asthma-relevant phenotypes between clusters 31 and 47. Asterisks indicate FDR<0.05.

**c**. Comparison of R-squared values of partitioned PRS models between EpiCop clusters vs baseline model.
